# HyperGenie: A method for predicting enzymatic gene essentiality using hypergraph neural networks and genome-scale metabolic models

**DOI:** 10.64898/2026.11.15.766798

**Authors:** Panayiotis Ioannou, Iulia Duta, Suraj Verma, Pietro Cicuta, Pietro Liò, Claudio Angione

**Affiliations:** Department of Physics, University of Cambridge, Cambridge, UK; Department of Computer Science and Technology, University of Cambridge, Cambridge, UK; School of Computing, Engineering and Digital Technologies, Teesside University, Middlesbrough, UK; Centre for Digital Innovation, Teesside University, Middlesbrough, UK; National Horizons Centre, Darlington, UK

**Keywords:** Gene essentiality, Metabolic modelling, Hypergraph neural networks, Deep learning

## Abstract

Identifying essential genes in bacteria is crucial for drug discovery and synthetic biology. Genome-scale metabolic models (GEMs) evaluated via flux balance analysis remain standard, yet they rely on the often-invalid premise that mutant strains optimize for maximum growth. Recent machine learning approaches circumvent this limitation using random forests or graph neural networks on mass flow graphs (MFGs). However, data-driven tools like random forests discard network topology, while graph-based tools like MFGs decompose multi-component biochemical reactions into pairwise interactions, misrepresenting complex stoichiometry and limiting interpretability. Furthermore, existing methods remain largely uncharacterised in data-scarce regimes and cross-environmental conditions.

Here, we introduce HyperGenie, a framework that integrates genome-scale metabolic models with directed hypergraph neural networks by representing metabolites as nodes and reactions as hyperedges. Operating directly on wild-type flux features, HyperGenie eliminates post-knockout optimality assumptions, achieves a 13% PR-AUC improvement over prior objective-dependent methods, and matches computationally demanding, state-of-the-art flux-sampling baselines. Using HyperGenie to guide targeted growth curve assays, we identified latent essentiality misannotations across three genes in the benchmark iML1515 dataset, which we then reconciled and validated experimentally via PCR.

In data-scarce regimes, HyperGenie retained robust predictive performance (> 90% PR-AUC) when trained on as little as 40% of the data. To isolate the value of hypergraph message passing, we benchmarked HyperGenie against an independent-reaction multilayer perceptron (MLPGenie) across diverse nutrient environments and uptake rates, where HyperGenie’s hypergraph inductive bias yielded consistent accuracy gains (> 2% in glucose; > 4% across six alternative carbon sources).

Notably, HyperGenie exhibits robust zero-shot cross-nutrient generalization, accurately forecasting gain-of-essentiality phenotypes on novel carbon sources, though loss-of-essentiality remains challenging. Finally, coupling hypergraph representations with SHAP attribution provides actionable mechanistic insights into metabolic rewiring, establishing a data-efficient, interpretable platform for discovering novel antibiotic targets.

## 1 Introduction

Essential genes are those required for bacterial viability [1, 2]. Identifying these genes, whose removal causes growth arrest or cell death, can accelerate advancements in drug discovery and synthetic biology. Experimental determination of gene essentiality, such as phenotypic screening, is resource-intensive, and essentiality varies with environmental context. Computational techniques for predicting gene essentiality have become more popular as a means to complement experimental efforts and accelerate the discovery of essential genes. The majority of these techniques use machine learning [3] or network-based methods [4] with data or features obtained from sequence information, protein-protein interaction (PPI) networks, and multi-omics data [5–17]. Genome-scale metabolic models (GEMs) [18, 19] can predict enzymatic gene essentiality [20] by linking metabolic reactions to genes via gene-protein-reaction (GPR) rules. Gene knockouts are simulated by disabling associated reactions, and constrained modelling methods like flux balance analysis (FBA) [21] predict their impact. FBA uses linear programming to optimise an objective, typically growth rate, under flux constraints. However, owing to the inherent redundancy of biological networks, the optimal solution identified by FBA is not unique; thousands of distinct flux distributions can achieve the same maximal objective value. Flux variability analysis (FVA) [22] captures this metabolic adaptability by calculating the minimum and maximum bounds of each reaction flux that can still sustain the optimal state. Therefore, the flux ranges calculated by FVA could serve as highly valuable, biologically grounded features for machine learning models to improve phenotypic predictions (e.g., gene essentiality).

Standard FBA on GEMs assumes that knockout strains optimise the same objective as the wild type, often growth, but this is not always accurate [23]. They can also misclassify essential genes, necessitating improved prediction methods. Recent approaches address these limitations by combining GEMs with machine learning, such as graph neural networks (FlowGAT) [24] and random forests [25]. These use FBA-derived fluxes to build mass flow graphs (MFGs) [26], representing the metabolic network as weighted, directed graphs with nodes as reactions and edges indicating metabolite flow. In this context, metabolic flux represents the rate at which molecules are transformed through biochemical reactions within a metabolic network, reflecting the activity level of different metabolic routes. To bypass the requirement of a predefined objective function and its inherent optimality assumption, newer architectures like FluxGAT employ flux sampling rather than relying on point estimates from FBA. This shift is particularly crucial for eukaryotic cells; while biomass optimization generally holds for bacteria [27, 28], it is usually inappropriate for mammalian cells [29]. Extending this paradigm, Flux Cone Learning (FCL) [30] has recently established state-of-the-art performance for predicting gene essentiality in the model bacterium *Escherichia coli* (*E.coli*). By training a random forest classifier on flux sampling data, FCL achieves predictive accuracies that surpass the standard FBA benchmark.

Despite these advances in flux generation and predictive accuracy, existing methods either discard explicit network topology (as in FCL) or rely on pairwise reaction-centric graphs like MFGs. Because MFGs project the network onto reaction nodes, they only indirectly represent metabolites and struggle to capture complex interactions involving multiple reactants and products. To address these representation challenges, hypergraphs offer a potential solution [31]. Hypergraphs are a generalization of standard graphs: while a typical graph uses edges to connect exactly two nodes, a hypergraph uses hyperedges that can connect any number of nodes simultaneously. This makes hypergraphs more suitable for modelling relationships where multiple entities interact at once, unlike graphs, which are limited to pairwise connections [32, 33].

Modelling bacterial metabolic networks as hypergraphs, with hyperedges representing reactions and nodes representing metabolites, offers several advantages over standard graph approaches like MFGs. Explicitly retaining metabolites as nodes preserves their central topological role in the network, facilitating the direct analysis of metabolite-centric properties alongside reaction fluxes. Furthermore, by utilizing directed and weighted hyperedges that partition reactants from products, hypergraphs faithfully encode reaction stoichiometry, directionality, and reversibility, providing a more comprehensive and detailed representation of metabolic networks [31, 34].

While hypergraph neural networks [35] have been successfully leveraged to map gene regulatory systems [36], can perform gap-filling in metabolic networks [37, 38], and predict growth phenotypes under diverse conditions [39], their potential for predicting gene essentiality within the framework of GEMs remains unexplored [40]. Leveraging this hypergraph representation, we introduce HyperGenie, a computational method for predicting enzymatic gene essentiality using hypergraph neural networks and GEMs.

In summary, this work makes the following contributions: **(1) A novel hypergraph architecture for metabolism:** We introduce HyperGenie, a method for predicting enzymatic gene essentiality using hypergraph neural networks and genome-scale metabolic models. By operating directly on wild-type flux distributions, HyperGenie circumvents the limiting assumption of Flux Balance Analysis (FBA) that knockout strains inherently optimize for maximum growth rate. **(2) State-of-the-art predictive performance:** HyperGenie achieves accuracy, precision, and F1-scores that match or exceed the FBA gold standard, establishing a highly competitive baseline alongside the current state-of-the-art method [30]. Notably, it yields a 13% PR-AUC improvement over prior objective-function-dependent approaches for *E. coli* grown aerobically on glucose [24, 25]. **(3) Value of hypergraph inductive biases over multi-layer perceptrons:** We evaluate the explicit contribution of hypergraph connectivity by comparing HyperGenie against an engineered multilayer perceptron baseline (MLPGenie) trained on identical constraint-based flux features. HyperGenie consistently outperforms this non-relational baseline, yielding a > 2% accuracy gain in glucose minimal medium and > 4% improvements across six alternative carbon sources. Crucially, beyond raw predictive gains, the hypergraph inductive bias provides topological context necessary for mechanistic and biological interpretability. **(4) High sample efficiency:** We establish the viability of both HyperGenie and MLPGenie in data-scarce regimes, demonstrating that robust predictive performance (PR-AUC > 90%) can be maintained using as little as 20% to 40% of the data for training. **(5) Cross-condition generalization and antibiotic target discovery:** We evaluate the models across diverse nutrient conditions and uptake rates. We demonstrate HyperGenie’s capacity for zero-shot generalization, training exclusively on glucose media and accurately predicting essentiality on alternative carbon sources, with a specific focus on identifying conditionally essential genes, highlighting its utility for discovering context-dependent antimicrobial targets. **(6) Discrepancy-driven discovery and experimental validation:** By systematically analysing predictions discordant with existing literature, we used HyperGenie to prioritize candidate genes for targeted growth curve assays. This model-guided approach uncovered and experimentally resolved latent essentiality misannotations across three genes in the widely used iML1515 dataset [19], demonstrating HyperGenie’s practical utility for data curation and novel phenotype discovery. **(7) SHAP-assisted biological interpretability:** To uncover the mechanistic basis of the model’s predictions, we conducted global and local feature attribution using SHapley Additive exPlanations (SHAP) [43]. Global SHAP summaries revealed that HyperGenie relies primarily on FVA bounds for its predictions. In addition, despite being trained exclusively on aerobic glucose M9 minimal medium, the model generalized to acetate M9 conditions to resolve key mechanistic adaptations: the directional reversal of central carbon flux between glycolysis and gluconeogenesis, carbon conservation via the glyoxylate shunt, and emergent biosynthetic vulnerabilities under two-carbon substrate limitation. At the local scale, Kernel SHAP demonstrated that gene essentiality is driven by multi-tier metabolic coupling, with key examples including the prioritization of immediate precursor supply, competitive downstream sinks of key intermediates, and distal modulators regulating shared cofactor pools.

## 2 Methodology

In this work, we represent the metabolic network as a hypergraph, with metabolites as nodes and reactions as hyperedges. To effectively capture higher-order relationships and the network’s complex topology, we leverage the recently proposed Hypergraph Neural Network (HGNN) [41]. Figure 1A illustrates the inputs to the HGNN. The HGNN operates on reaction-level input features derived from FBA, parsimonious FBA (pFBA) and FVA. Crucially, while these features are assigned to individual reactions, they are calculated over the entire network and inherently encapsulate global, systems-level metabolic dependencies. Finally, for each reaction (hyperedge) that has a many-to-one or one-to-one mapping with a gene, we generate a binary essentiality prediction for its corresponding gene using gene-protein-reaction (GPR) rules. In the many-to-one scenario, the gene’s essentiality is determined by the majority vote of the essentiality predictions of the reactions linked to that gene.

**Fig. 1:**
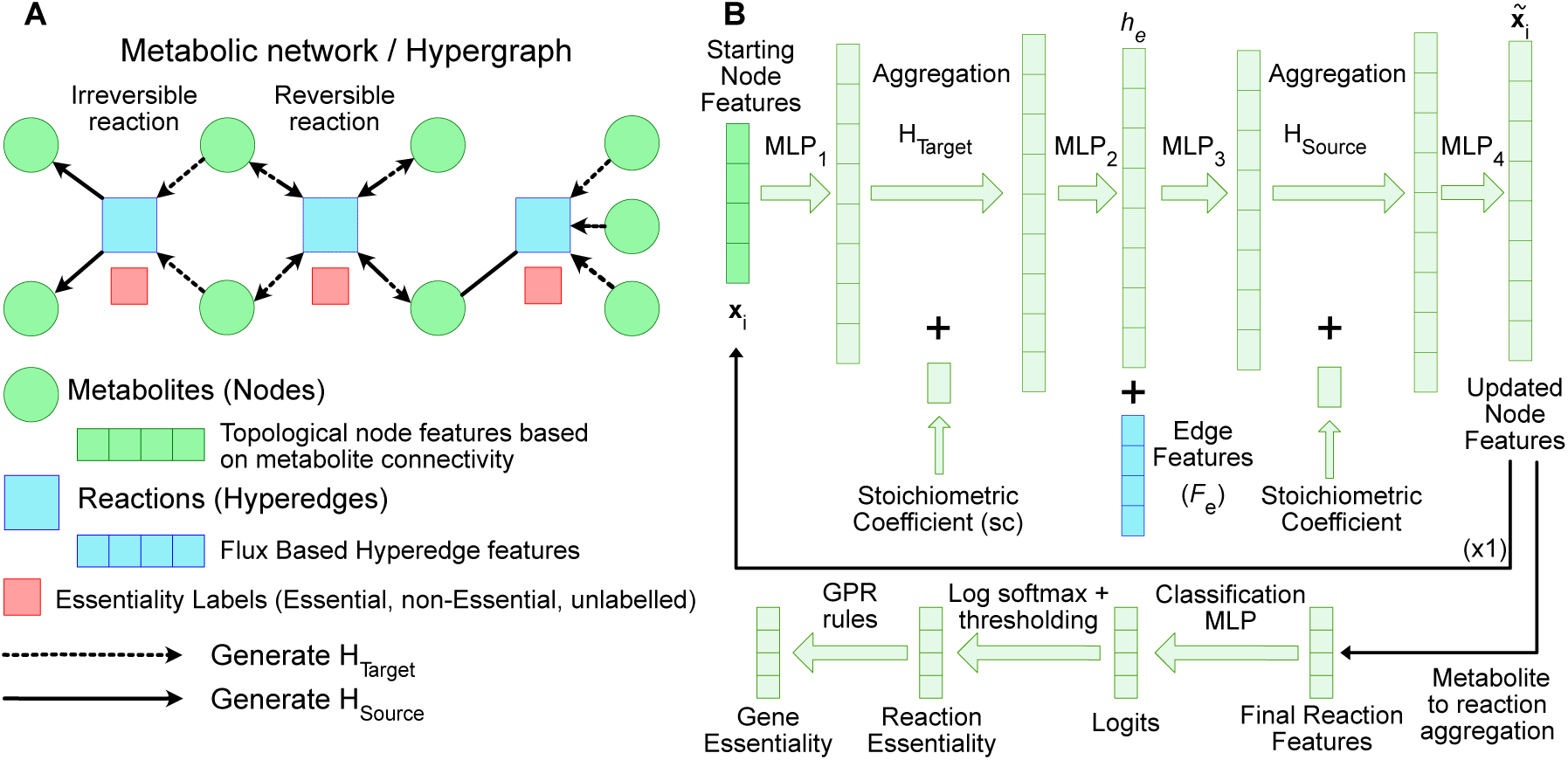
Overview of the directed hypergraph neural network framework for predicting enzymatic gene essentiality. To capture higher-order biochemical interactions beyond pairwise graph representations, our framework models metabolic networks as directed hypergraphs and integrates metabolite- and reaction-level features via a hypergraph neural network to predict essential genes. **(A) Schematic hypergraph neural network input**, showing a small hypothetical metabolic network where metabolites (nodes) are connected by directed biochemical reactions (hyperedges). **(B) Architectural schematic of the hypergraph neural network layer deployed in this work.** We adapted the AllDeepSets framework [41] to reflect metabolic network structure by incorporating asymmetric incidence matrices (**H**_Source_, **H**_Target_), stoichiometric coefficients, and reaction-level flux features. From left to right: MLP_1_ first projects initial metabolite features (**x***_i_*) into latent representations, where each **x***_i_* is constructed by concatenating multi-step incoming and outgoing topological flow profiles adapted for hypergraphs from Cooper and Barahona [42] (Section 2.1.1). Before aggregating product metabolites into reactions via the target matrix (**H**_Target_), stoichiometric coefficients are concatenated (+) to the node embeddings. Following aggregation, MLP_2_ computes intermediate reaction embeddings (**h***_e_*), which are concatenated (+) with reaction features (**F***_e_*) and refined via MLP_3_. For the reverse message passing back to reactant metabolites, hyperedge features and stoichiometric coefficients are aggregated using the source matrix (**H**_Source_). Subsequently, MLP_4_ processes the pooled representations to produce the updated metabolite embeddings. After the complete model performs two message-passing iterations through this layer, the resulting metabolite features are aggregated according to reaction connectivity to construct the final reaction (hyperedge) embeddings. These reaction representations are then passed through an MLP classification head followed by a log-softmax activation to compute class log-probabilities. Finally, each reaction is classified as essential or non-essential via probability thresholding or argmax selection, and reaction essentiality is mapped to gene essentiality using gene–protein–reaction (GPR) associations.

To isolate the specific contribution of hypergraph-based message passing, we also implemented a multilayer perceptron baseline, MLPGenie, to benchmark against our method. MLPGenie shares the same input features, architecture scale, and GPR mapping pipeline as HyperGenie, but completely omits the hypergraph propagation layers, evaluating each reaction as topologically independent during training. However, because it relies on the same FBA, pFBA and FVA input features, MLPGenie still retains implicit information about the global metabolic state.

### 2.1 Hypergraph construction

We construct a hypergraph representation based on the iML1515 model [19], the most comprehensive genome-scale reconstruction of *E. coli* K-12 MG1655 metabolism currently available, and its associated stoichiometric matrix. This model, obtained from the BiGG model database [44], encompasses 2712 reactions, 1877 metabolites, and 1516 enzymatic genes (as of December 2024).

Unlike standard directed graphs that only link pairs of nodes, a directed hypergraph uses directed hyperedges to natively connect multiple source nodes (reactants) to multiple target nodes (products) simultaneously [31–33]. In our hypergraph, metabolites are represented as nodes and reactions as hyperedges, naturally preserving the multi-component structure and inherent directionality of biochemical reactions. Irreversible reactions are modelled as directed hyperedges to reflect one-way mass flow, whereas reversible reactions are represented as undirected hyperedges to accommodate bidirectionality. Within our computational model, feature aggregation propagates opposite to the natural biochemical flux, passing information backward from products to reactants. This inverse message passing is crucial because, while physical mass flows forward, biological context and downstream demand flow backward: this allows upstream metabolites to encode downstream pathway dependencies, cellular demand, and final metabolic outcomes.

To precisely define the hypergraph structure, we utilise source and target incidence matrices. These matrices are derived from a binary stoichiometric matrix, where entries indicate the presence (1) or absence (0) of a metabolite in a reaction, regardless of the specific stoichiometric coefficient. The source incidence matrix, **H***_source_*, identifies the reactants consumed in each reaction, while the target incidence matrix, **H***_target_*, identifies the products generated (Figure 1A).

To integrate genes and their essentiality into the hypergraph, we use GPR rules to link reactions (hyperedges in the case of the hypergraph) to their corresponding genes. These rules specify the connection between genes, their encoded enzymes, and the biochemical reactions the enzymes catalyse. Since our method makes predictions at the hyperedge (reaction) level, we map gene essentiality labels to reactions using the GPR rules to produce reaction essentiality labels. This mapping is feasible only when genes have a one-to-one or oneto-many relationship with reactions, allowing a direct conversion of gene essentiality to reaction essentiality, a strategy first used by Hasibi et al. [24].

To align our analysis with the BW25113 strain, commonly used in gene essentiality experiments, we adjust the iML1515 model based on GPR rules to reflect its genetic differences from the K-12 MG1655 strain. This adjustment demonstrates how our model encodes genetic variations. Specifically, we set the lower and upper flux bounds of reactions associated with the *araBAD* and *rhaBAD* operons, and the *lacZ* gene to zero (Table D1). This simulates gene knockouts by restricting metabolite flow through the affected reactions, mimicking the absence of the corresponding enzymes. These modifications directly alter the hypergraph’s constraints, reflecting the strain-specific metabolic capabilities. Our model accommodates genetic differences either by uploading a strain-specific metabolic network or by modifying an existing one using GPR rules.

Using the resulting iML1515 model, we perform parsimonious flux balance analysis (pFBA) under aerobic conditions for different carbon sources; exact nutrient details are provided in Table D1. For glucose, this analysis identified 442 reactions with non-zero flux. Reactions with zero flux were deemed non-essential, as they are not required for growth under the specific condition. Of these 442 active reactions, 174 exhibit a one to-one relationship with a single gene, while 80 have a many-to-one gene association. This results in a total of 254 labelled reactions, which were used to train and assess HyperGenie. The case of one-to-many gene-to-reaction mappings is more complex. Although we include these for training purposes to expand the training dataset and to be able to make direct comparisons with Hasibi et al. [24], it is important to acknowledge the assumption that if a gene is essential, it does not necessarily imply that all reactions catalysed by that gene are also essential.

#### 2.1.1 Node features

Due to the lack of directly relevant metabolite information that could be utilised as features, we modified a feature generation method originally proposed by Cooper and Barahona [42] to generate topological features suitable for HGNNs. Specifically, Equations 1 and 2 were used to generate a feature vector for each node. In these equations, the vector **x***_i_* represents the node features of node *i*, the scalar values *a* and *o* represent the number of nodes that are connected to node *i* via incoming and outgoing paths of length *k*, respectively, *n* is the number of hyperedges in the hypergraph, **1***^n×^*^1^ is an *n*-dimensional vector of ones, **H***^T^_source_* is the transpose of the target incidence matrix, **H***_source_* is the source incidence matrix and [*· · ·*]*^k^* represents the matrix to the power of *k*.

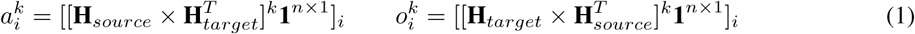

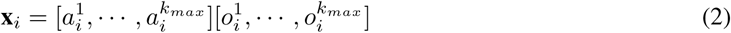

For each node *i*, Equation 1 is used to determine the number of nodes connected to it through incoming or outgoing paths of a specific length *k*. This is performed for *k* values ranging from 1 to a predefined maximum, *k_max_*(set to 4 in this work). Equation 2 describes the process of concatenating the counts obtained from Equation 1 for each *k*, creating the complete feature vector for node *i*.^1^

#### 2.1.2 Hyperedge features

Since hyperedges represent reactions, any flux-based features associated with each reaction are highly informative. We employ mathematical methods to simulate bacterial metabolism using genome-scale metabolic models, extracting flux-based hyperedge features from the results of three approaches: FBA, pFBA, and flux variability analysis (FVA) using a pFBA factor of 1.1 and a fraction of optimum of 0.99. The final hyperedge features are obtained by concatenating the outcomes of the above results. Further mathematical details regarding FBA and FVA are provided in Appendix A.

#### 2.1.3 Model architecture

The hypergraph neural network architecture developed in this study builds upon the AllDeepSets framework [41], introducing two key biological and structural adaptations. First, we incorporate reaction-specific edge features directly into the hyperedge embedding space. Because hyperedges correspond to metabolic reactions, integrating flux-based features provides a biophysically principled representation of pathway activity, enabling the network to resolve metabolic bottlenecks, alternative routing, and reaction reversibility. Second, we integrate stoichiometric coefficients directly into the message-passing propagation layers. This enables the model to explicitly encode net molar turnover and thermodynamic constraints governing the metabolic network, ultimately improving gene essentiality prediction.

Formally, our modified AllDeepSets message-passing layer is defined in Equation 3. Here, MLP represents a multilayer perceptron, *ν* signifies the set of nodes, *ε* the set of hyperedges, *sc_i,e_* the stoichiometric coefficient of node (metabolite) *i* within hyperedge (reaction) *e*, *h_e_*the hyperedge embedding, *∗ → e* represents the source nodes of hyperedge *e*, *∗ → i* represents all the hyperedges that have node *i* as a target, *F_e_* represents the hyper-edge features and **x***_i_* represents the updated features of node *i*. The model hyperparameters can be found in Table D2.

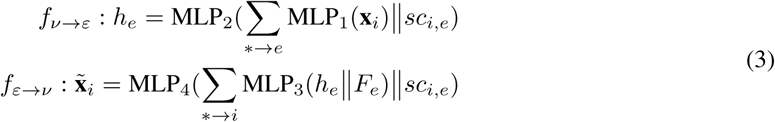

Figure 1B shows the details of the HyperGenie method. The initial feature vectors of each node (**x**) in the hypergraph are first embedded using an MLP. Then, we perform an aggregation on the transformed node features using the target incidence matrix, **H***_target_*. During this aggregation, we concatenate a coefficient (*sc_i,e_*) to each node-hyperedge pair, corresponding to the stoichiometric coefficient of each metabolite within a given reaction. The aggregated stoichiometric coefficients incorporate information on the molar changes that occur with each reaction. Next, another MLP is applied to the aggregated features, generating a set of hyperedge-specific features (*h_e_*). This sequence of operations constitutes the *f_ν→ε_* part of Equation 3.

The resulting features are then concatenated with the flux-based hyperedge features (*F_e_*), and the above process is repeated, but with the aggregation operation now guided by the source incidence matrix, **H***_source_*. This reversed aggregation updates the node features based on information propagated from the hyperedges (see the *f_ε→ν_* part of Equation 3). After the last HGNN layer is applied, we employ summation readout to derive the final hyperedge features. This involves summing the updated node features (**x***_i_*) for all nodes within each hyperedge, regardless of direction. Finally, an MLP classifier produces the final hyperedge embeddings.

### 2.2 MLPGenie

For MLPGenie, each input node represents a distinct metabolic reaction. Mirroring our approach for the features of the hypergraph edges (Section 2.1.2), we assigned flux-based features to each reaction. We trained and assessed this method using the 254 labelled reactions detailed in Section 2.1.

The MLPGenie architecture comprises two fully connected layers, structurally mirroring the classification module of HyperGenie. The training procedure was similarly standardized against the hypergraph model, utilizing the same optimization parameters, learning rate, and maximum epochs. The sole deviation was the weight decay, which was set to zero for MLPGenie. A comprehensive summary of the evaluated search space and final selected hyperparameters for both models is provided in Table D2.

### 2.3 Dataset

The models (HyperGenie and MLPGenie) were trained using gene essentiality data obtained from Monk et al. [19]. These data characterise the growth of single-gene knockouts of the *E. coli* BW25113 strain in minimal media across 16 different carbon sources. Genes were classified as essential if their deletion resulted in a growth rate below 0.5; otherwise, they were deemed non-essential. Of the 1516 enzymatic genes in the iML1515 model, experimental essentiality data were available for 1503 genes for each carbon source, with the exception of mannitol, for which data were available for only 1072 genes.

HyperGenie, a binary classifier employing a semi-supervised learning approach, is trained on 254 reaction essentiality labels (193 essential and 61 non-essential) derived from pFBA analysis, as detailed in Section 2.1. Crucially, the remaining reactions (hyperedges) in the hypergraph are included as unlabelled input to the model. This allows HyperGenie to exploit the underlying network structure and relationships between reactions and metabolites, even in the absence of explicit labels. During training, both HyperGenie and MLPGenie were optimized by minimizing the negative log-likelihood loss. To indicate specific feature preprocessing strategies, models utilizing log-modulus normalization of hyperedge features are denoted with the suffix “Norm” (e.g., HyperGenie Norm). Consequently, unless explicitly stated otherwise, all general references to HyperGenie throughout the text denote the standard, unnormalized model.

## 3 Results

Beyond comparing with top methods in the literature, we systematically evaluate the predictive capabilities, robustness, and interpretability of HyperGenie and MLPGenie. First, we optimize metabolic flux features and assess model robustness under data-scarce regimes by training on small subsets of the available data, demonstrating high sample efficiency. To determine environmental adaptability, we test the models across varying nutrient conditions and flux uptake rates, revealing their capacity for zero-shot generalization across diverse environments. Furthermore, through failure-mode analysis, we identify and correct misannotations within the experimental dataset. Finally, we apply SHAP to provide mechanistic interpretability for the predictions.

### 3.1 Performance evaluations

To evaluate the performance of our models, we conducted a 5-fold cross-validation process 10 times, using different folds and random seeds for a total of 50 runs. Under identical splits and conditions, both HyperGenie and MLPGenie outperformed FlowGAT [24] and Random Forest [25] by over 12% in PR-AUC (Table 1). Other methods mentioned in the introduction are not directly comparable due to differences in training data, experimental conditions, and cell or bacterial types. Since the FCL method [30] generates predictions directly at the gene level, a direct comparison using the PR-AUC metric is not possible; however, a comparison using other performance metrics, such as accuracy, is provided in Section 3.1.4. Figure D1 shows precision-recall curves for one of the 5-fold cross-validation cycles. We note that, for this particular evaluation setup, the application of log-mod normalization to the edge features did not impact the PR-AUC results.

**Table 1:** Gene essentiality prediction performance in *E. coli* (aerobic M9 glucose medium). Performance is evaluated using PR-AUC (%), reported as mean ± standard deviation across 50 runs with 5-fold cross-validation, and compared against literature baselines. Sections compare the proposed models using only FBA as an edge feature versus the complete feature set, followed by an ablation study on hypergraph message-passing directionality (where the default model employs backward propagation). Bold values indicate the highest overall performance.

| **Model Architecture** | **PR-AUC (%)** |
|---|---|
| *Literature Baselines* |  |
| FlowGAT [24] | 82.5 |
| Random Forest [25] | 83.0 |
| *FBA-Only Feature (100 runs)* |  |
| MLPGenie | 84.4 ± 7.2 |
| HyperGenie (Ours) | 87.3 ± 7.3 |
| *Full Feature Set* |  |
| HyperGenie (Ours) | **96.1 ± 2.6** |
| HyperGenie Norm (Ours) | 96.0 ± 2.3 |
| MLPGenie | 95.8 ± 2.8 |
| *Graph Directionality Ablations* |  |
| HyperGenie Forward (Ours) | 90.6 ± 4.3 |
| HyperGenie No Direction (Ours) | 93.4 ± 3.3 |

#### 3.1.1 Hypergraph directionality

Crucially, the directionality of message passing within the hypergraph architecture played a significant role in overall model performance (Table 1). While it is intuitive to assume that aligning hypergraph propagation with the biological direction of metabolic reactions would yield optimal results, our empirical findings demonstrate the inverse: reversing the propagation direction resulted in a performance improvement of greater than 5% in PR-AUC.

When hypergraph propagation is reversed compared to the natural flow of reactions, the model learns more effectively. Reverse message passing enables each substrate to aggregate information from all subsequent reactions and products, effectively allowing the architecture to learn the downstream biological demand that drives metabolic activity. This approach naturally highlights metabolic bottlenecks and crucial pathways that act as the sole suppliers of critical biomass precursors. Consequently, the model can effectively map biological dependencies, recognizing that severing a single upstream reaction will disrupt everything downstream.

#### 3.1.2 The importance of flux-based edge features for predicting gene essentiality

To enhance the predictive performance of our model, we investigated the impact of incorporating metabolic flux information as hyperedge features. Specifically, we considered four distinct types of flux data derived from the metabolic network: FBA, pFBA, and the minimum and maximum values from flux variability analysis (FVA). We evaluated different combinations of these flux features to determine which combination yielded the best performance in predicting gene essentiality. The results are summarised in Figure 2A. The best results were obtained when using all four flux types or when both FVA fluxes were combined with either FBA or pFBA. Mann-Whitney U tests show no significant difference between these 3 combinations but significant differences with all other combinations.

**Fig. 2:**
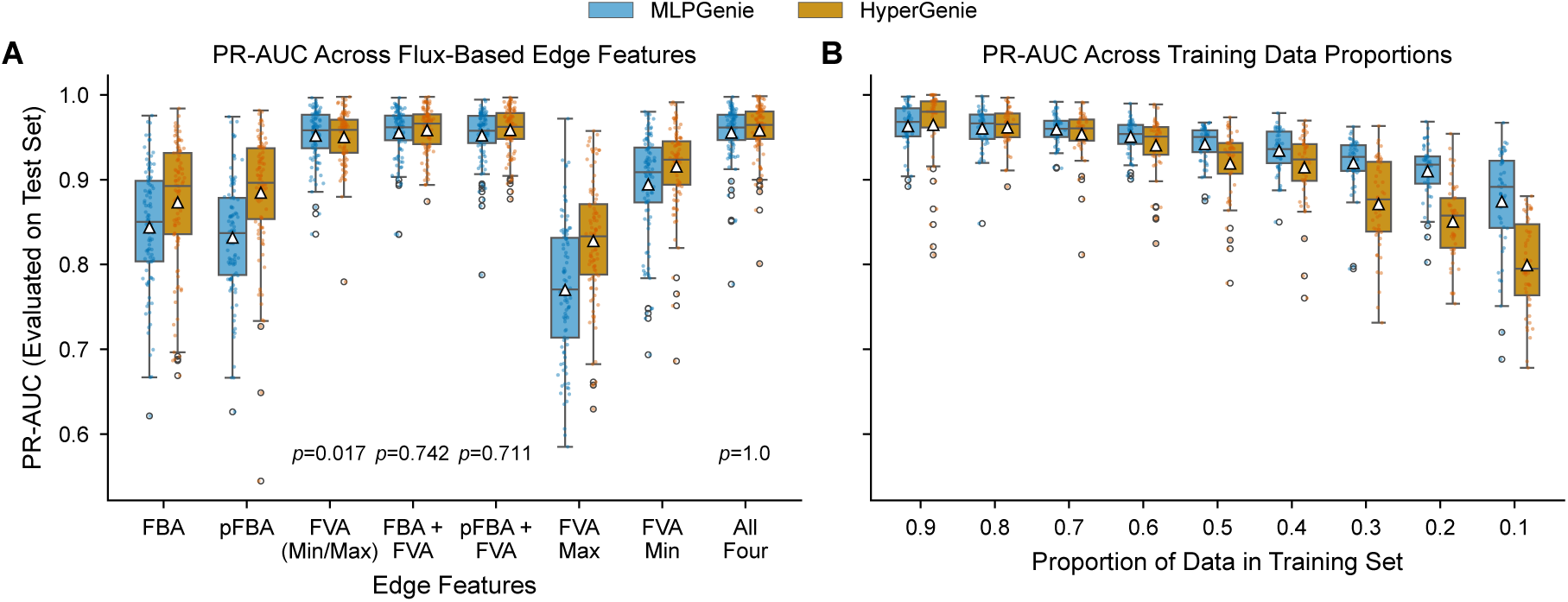
Flux Variability Analysis (FVA) bounds drive peak gene essentiality prediction and enable high performance even with limited training data. (A) Combining both FVA bounds with either FBA or pFBA achieves optimal predictive performance, matching the complete four-feature set. PR-AUC scores are shown across various combinations of FBA, pFBA, and FVA (min/max) flux features for MLPGenie (blue, left) and HyperGenie (orange, right). Triangles indicate mean scores over 100 runs. Mann-Whitney U tests compare each combination against *All Four* (*p <* 0.05 considered significant; *p <* 0.01 not shown). (B) Predictive performance remains robust with PR-AUC exceeding 90% even when training data is reduced to 40%. PR-AUC scores across varying training set sizes over 50 random runs. Below the 50% training threshold, HyperGenie performance degrades more rapidly than MLPGenie due to sample size constraints. All results are for *E. coli* grown aerobically in glucose minimal media.

The minimum flux boundary derived from FVA proved most informative for predicting gene essentiality. Mathematically, a minimum flux of zero proves that alternative, redundant pathways exist, while a non-zero minimum indicates that a reaction is obligatory. Conversely, the maximum flux feature yielded the lowest performance when used alone. This is expected, as maximum flux merely defines a reaction’s potential capacity. Capacity alone cannot determine essentiality; identifying a true metabolic bottleneck requires knowing the necessary downstream demand, not just the maximum possible throughput. In contrast, FBA and pFBA solutions, despite offering insights into optimal flux distributions, are not unique.

When utilizing the complete four-feature set, the PR-AUC difference between HyperGenie and MLPGenie was not statistically significant (Mann-Whitney U test, *p* = 0.354). Conversely, when relying exclusively on FBA fluxes, HyperGenie maintained a statistically significant performance advantage of approximately 3% over MLPGenie (Table 1, *p* = 0.0014). This similar performance when using the complete feature set is initially counter-intuitive, given the capacity of hypergraphs to natively map the higher-order, non-linear topological interactions inherent to metabolic reactions. However, we hypothesize that FVA bounds act as a proxy for the network’s global structure. This gives MLPGenie enough information about the network’s layout to perform just as well as the hypergraph architecture in terms of the PR-AUC performance metric.

Similar to previous works [24, 25], our method uses fluxes derived from the wild-type strain. Predicting gene essentiality through FBA approaches follows the assumption that knockout strains optimise for growth rate. However, this may not always be valid, as gene knockouts can disrupt cellular processes and lead to suboptimal or alternative growth strategies.

#### 3.1.3 Robustness to limited training data

Figure 2B displays the PR-AUC values, calculated on the test sets, obtained from multiple model runs with varying training proportions. These results were generated by randomly dividing the dataset into training and test sets 50 times. Both HyperGenie and MLPGenie efficiently predict gene essentiality, achieving a PR-AUC exceeding 90% with only 40% of the data used for training. However, when the training proportion drops below 50%, HyperGenie’s performance falls off much faster. Because the hypergraph model is highly complex and has many more parameters, it is more prone to underfitting than the simpler MLP when data is severely limited. Despite this drop, these findings highlight the potential of both models to complement experimental studies by providing robust predictions even with small amounts of training data.

#### 3.1.4 Gene essentiality prediction: outperforming iML1515

We evaluated HyperGenie’s capacity for gene essentiality prediction against prior literature methods and the gold standard of FBA on the iML1515 genome-scale metabolic model (Table 2). For *E. coli* simulated under aerobic growth in glucose-supplemented M9 minimal media, HyperGenie demonstrates superior predictive performance compared to FBA of the iML1515 genome-scale metabolic model [19], FlowGAT [24], and Random Forest [25]. Furthermore, HyperGenie matches the current state-of-the-art FCL method [30] across two evaluable datasets of 136 and 137 genes. Because FCL was trained and evaluated using random splits, the published data naturally omits predictions for genes that did not happen to be included in the test sets. Additionally, HyperGenie’s current architecture is restricted to predicting essentiality for genes characterized by one-to-one or one-to-many gene-protein-reaction (GPR) rules. As a result, the comparison is limited to the overlapping subset of genes resolvable by both methods. Finally, HyperGenie outperforms the MLPGenie architecture by more than 2% in accuracy and more than 1% in F1 score.

**Table 2:**
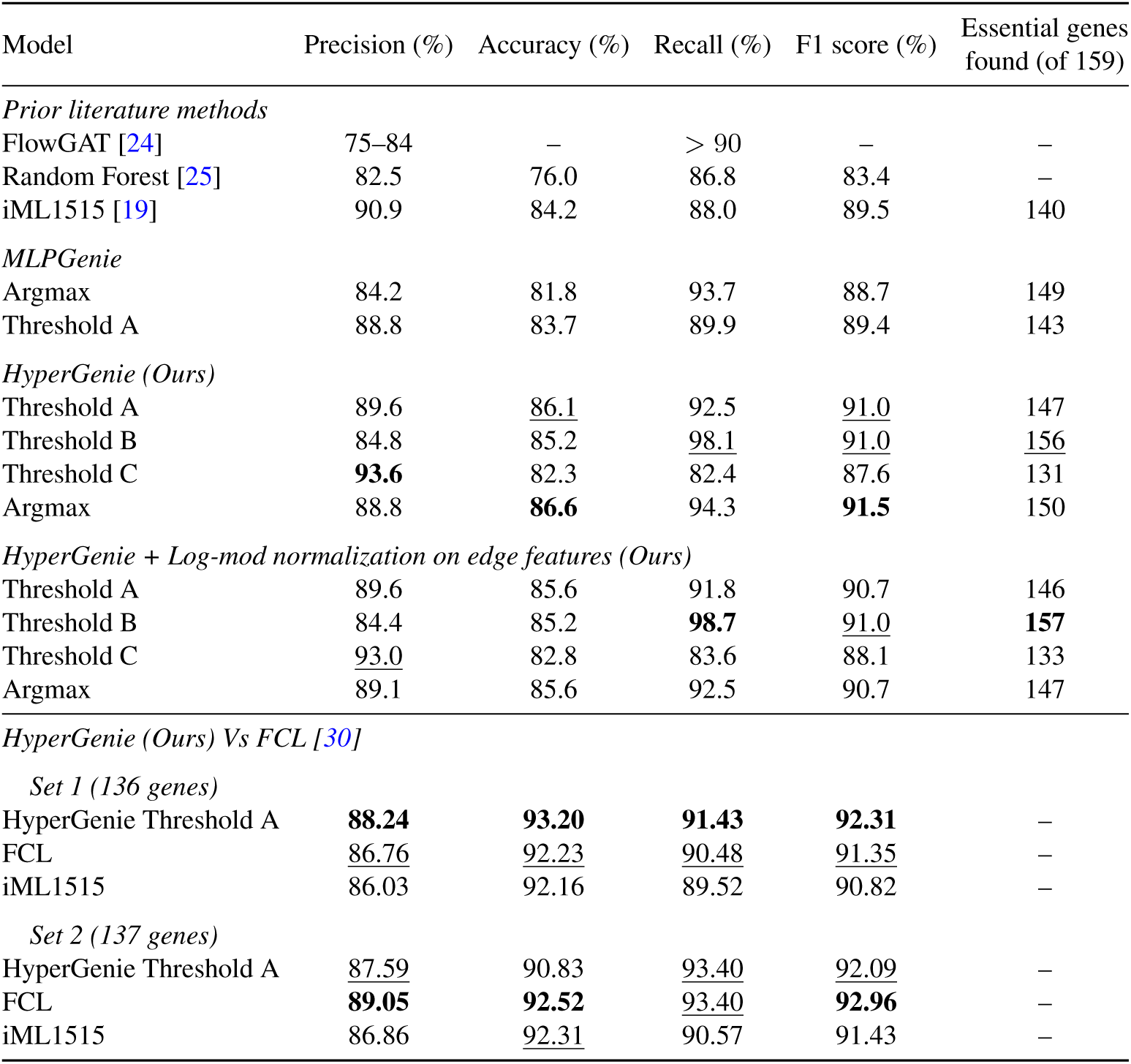
HyperGenie outperforms iML1515 mechanistic predictions and matches the state-of-the-art gene essentiality model (FCL). Performance was evaluated for *E. coli* in aerobic, glucose-supplemented M9 minimal media. For HyperGenie and MLPGenie, reaction predictions were mapped to gene essentiality via gene-protein-reaction (GPR) rules; final labels reflect a majority vote across 100 independent runs (ties defaulted to non-essential). “Argmax” denotes standard binary classification from output logits. Thresholds A, B, and C represent dynamic boundaries on the precision-recall curve classifying the top *n*, *n* + 5, and *n −* 5 ranked reactions as essential (*n* = true number of essential reactions). These thresholds optimize for downstream objectives like recall-driven metabolic engineering (B) or precision-driven drug discovery (C). The table’s first section compares MLPGenie and prior literature to HyperGenie (evaluated both with and without edge feature normalization), while the second compares HyperGenie to FCL on shared subsets of gene predictions. Bold and underlined values denote the best and second-best scores, respectively.

| Model | Precision (%) | Accuracy (%) | Recall (%) | F1 score (%) | Essential genes found (of 159) |
|---|---|---|---|---|---|
| *Prior literature methods* |  |  |  |  |  |
| FlowGAT [24] | 75–84 | – | > 90 | – | – |
| Random Forest [25] | 82.5 | 76.0 | 86.8 | 83.4 | – |
| iML1515 [19] | 90.9 | 84.2 | 88.0 | 89.5 | 140 |
| *MLPGenie* |  |  |  |  |  |
| Argmax | 84.2 | 81.8 | 93.7 | 88.7 | 149 |
| Threshold A | 88.8 | 83.7 | 89.9 | 89.4 | 143 |
| *HyperGenie (Ours)* |  |  |  |  |  |
| Threshold A | 89.6 | 86.1 | 92.5 | 91.0 | 147 |
| Threshold B | 84.8 | 85.2 | 98.1 | 91.0 | 156 |
| Threshold C | **93.6** | 82.3 | 82.4 | 87.6 | 131 |
| Argmax | 88.8 | **86.6** | 94.3 | **91.5** | 150 |
| *HyperGenie + Log-mod normalization on edge features (Ours)* |  |  |  |  |  |
| Threshold A | 89.6 | 85.6 | 91.8 | 90.7 | 146 |
| Threshold B | 84.4 | 85.2 | **98.7** | 91.0 | **157** |
| Threshold C | 93.0 | 82.8 | 83.6 | 88.1 | 133 |
| Argmax | 89.1 | 85.6 | 92.5 | 90.7 | 147 |
| *HyperGenie (Ours) Vs FCL [30]* |  |  |  |  |  |
| *Set 1 (136 genes)* |  |  |  |  |  |
| HyperGenie Threshold A | **88.24** | **93.20** | **91.43** | **92.31** | – |
| FCL | 86.76 | 92.23 | 90.48 | 91.35 | – |
| iML1515 | 86.03 | 92.16 | 89.52 | 90.82 | – |
| *Set 2 (137 genes)* |  |  |  |  |  |
| HyperGenie Threshold A | 87.59 | 90.83 | 93.40 | 92.09 | – |
| FCL | **89.05** | **92.52** | 93.40 | **92.96** | – |
| iML1515 | 86.86 | 92.31 | 90.57 | 91.43 | – |

A key advantage of HyperGenie is its capacity for calibrated thresholding along the precision-recall curve, allowing decision boundaries to be tailored to distinct biological objectives. To illustrate this flexibility, we defined three operating thresholds based on the true number of essential reactions in the test set (*n*). Threshold A classifies the top *n* ranked reactions as essential, whereas Thresholds B and C classify the top *n* + 5 and *n −* 5 reactions, respectively. This tunability enables task-specific optimization. For example, in drug target discovery, the conservative decision boundary of Threshold C maximizes precision to minimize potential off-target effects. Conversely, for metabolic engineering applications, the relaxed boundary of Threshold B prioritizes recall, capturing a broader set of essential genes necessary for robust biochemical production. Standard binary classification, derived by applying the argmax function directly to the output logits, similarly yields highly robust baseline predictions. Finally, HyperGenie’s predictive performance remains stable when log-mod normalization is applied to the network edge features.

Gene essentiality predictions (Table 2) were derived by translating HyperGenie’s reaction essentiality labels via GPR rules. Results from 50 runs were aggregated, with genes assigned a final label by majority vote; ties defaulted to non-essential.

#### 3.1.5 Discrepancy analysis exposes dataset annotation errors and uncharacterised metabolic bypasses

We systematically evaluated persistent discrepancies between predictions from the machine learning architectures (MLPGenie, as well as HyperGenie with and without log-mod normalization) and the empirical labels in the iML1515 database [19]. To ensure robust evaluation, we performed 200 iterations of 5-fold cross-validation (1,000 total training runs), yielding 200 independent predictions for each reaction. Reaction essentiality outputs were subsequently mapped to gene essentialities using Gene-Protein-Reaction (GPR) rules (Table 3). In Table 3, low agreement scores highlight frequent conflicts with the database, revealing two distinct categories of discrepancy: erroneous pathway-level essentiality annotations in the iML1515 dataset and unannotated metabolic bypasses in the reconstruction.

**Table 3:** Persistent model–database discrepancies across 200 independent evaluations. Values represent the frequency (out of 200 runs) with which the models’ predictions matched the benchmark iML1515 empirical labels [19]. Low values highlight consistent disagreement with the database, exposing annotation errors in pantothenate (B_5_) and biotin (B_7_) biosynthesis. The first section of the table shows the top 5 most incorrect genes, while the second section shows relevant genes.

| **HyperGenie (HG)** |  |  |  | **MLPGenie** |  |  |
|---|---|---|---|---|---|---|
| Gene | HG | HG Norm | Total Runs | Gene | MLPGenie | Total Runs |
| *Top 5* |  |  |  |  |  |  |
| *panC* (b0133) | 0 | 0 | 200 | *panD* | 0 | 200 |
| *hemF* (b2436) | 0 | 0 | 200 | *panC* | 0 | 200 |
| *kdsC* (b3198) | 0 | 0 | 200 | *panB* | 0 | 200 |
| *ubiH* (b2907) | 1 | 1 | 200 | *kdsC* | 0 | 200 |
| *mdh* (b3236) | 3 | 3 | 200 | *mdh* | 0 | 200 |
| *Relevant Genes* |  |  |  |  |  |  |
| *panD* (b0131) | 44 | 63 | 200 | *bioA* | 5 | 200 |
| *panB* (b0134) | 6 | 21 | 200 | *bioB* | 197 | 200 |
| *bioA* (b0774) | 58 | 21 | 200 | *bioF* | 7 | 200 |
| *bioB* (b0775) | 172 | 172 | 200 | *bioC* | 12 | 200 |
| *bioF* (b0776) | 162 | 170 | 200 | *bioD* | 10 | 200 |
| *bioC* (b0777) | 122 | 110 | 200 | *bioH* | 5 | 200 |
| *bioD* (b0778) | 45 | 70 | 200 |  |  |  |
| *bioH* (b3412) | 45 | 45 | 200 |  |  |  |

The most pronounced discrepancy occurred within the pantothenate (vitamin B_5_) and biotin (vitamin B_7_) pathways. Under unsupplemented glucose M9 minimal conditions, pantothenate biosynthesis is obligate [45]; yet, the iML1515 empirical dataset records *panB*, *panC*, and *panD* as non-essential. All evaluated models robustly overrode the database label to predict their biological essentiality. Similarly, within the biotin pathway, iML1515 designates only *bioB* as essential while marking the remaining five genes (*bioACDFH*) as non-essential. Because biotin synthesis cannot be bypassed and unsupplemented minimal medium lacks an exogenous source, this entire pathway is conditionally essential [45, 46].

Remarkably, despite noisy training labels, the models frequently recovered the true biological phenotype. However, for the HyperGenie models, *bioF* and *bioC* were misclassified relative to the actual empirical essentiality (Table 3). Because HyperGenie explicitly incorporates network topology, internally conflicting training signals, such as labelling *bioB* as essential while its pathway partners are marked non-essential, can propagate contradictory constraints through the hypergraph. In contrast, MLPGenie processes reaction features independently without topological priors, making it less susceptible to localized structural label contradictions and allowing it to identify the correct essentiality state.

In contrast to annotation errors, the failure mode observed for *kdsC* (b3198) stems from incomplete biological knowledge in the metabolic reconstruction. Here, the model’s false-essential prediction can be attributed to uncharacterised bypass mechanisms, as Sperandeo et al. [47] demonstrated that alternative proteins can functionally rescue the loss of *kdsC*.

To identify which genes disproportionately degrade model performance, we analysed the frequency of gene presence in the test set during cross-validation folds where HyperGenie achieved a PR-AUC below 0.92 (Table 4). The *kdsC* gene is overwhelmingly overrepresented in these low-performing folds, appearing twice as frequently as the next most common gene. For comparison, we also tracked gene frequencies in highperforming folds (PR-AUC > 0.96), as shown in Table 4. Together, these analyses indicate that incorrect predictions for *kdsC*, *hemF*, and *panC* exert a significantly harsher penalty on the overall PR-AUC metric than missclassifications of other genes. Representative PR-AUC curves illustrating these low-performing runs are provided in Figures D2A and D2B.

**Table 4:**
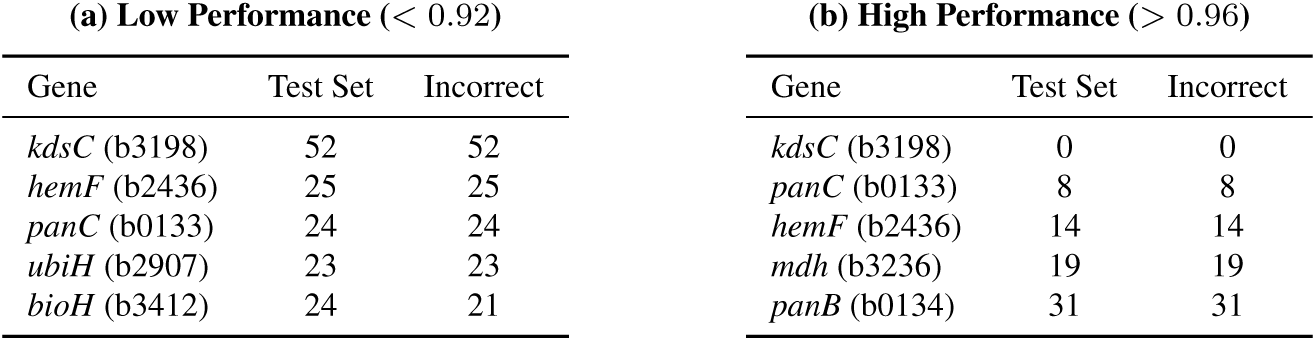
Impact of specific gene misclassifications on PR-AUC. The tables contrast the frequency of specific genes in the test set during: **(a)** low-performing folds, versus **(b)** high-performing folds. Genes that appear frequently in (a) but rarely in (b), such as *b3198*, exert a disproportionately severe penalty on the over-all PR-AUC metric when misclassified.

Table 5 details the predictive performance of HyperGenie under two targeted manipulation conditions: removing specific gene sets from the evaluation (Condition 1) and correcting their ground-truth essentiality labels to reflect biological truth (Condition 2). Set A comprises the genes explicitly identified as incorrectly annotated in the iML1515 dataset. The subsequent sets expand upon this baseline by incorporating genes such as *kdsC* and *mdh* to quantify the exact penalty these high-impact genes exert on the model. Removing Set A from the analysis (Condition A1) increases the PR-AUC to 96.7%, while correcting its labels (Condition A2) yields a comparable improvement to 96.6%. Notably, removing the Set B genes increases the PR-AUC even further (98.1%), which supports our previous finding that these genes (*kdsC* and *mdh*) disproportionately affect model performance.

**Table 5:** Impact of gene set ablation and label correction on PR-AUC. The table defines specific subsets of misclassified genes (**a**) and evaluates model performance across various manipulation conditions (**b**). Condition **1** corresponds to removing the reactions associated with the specified gene set, while Condition **2** modifies the true essentiality data for that set to reflect biological reality.

| **(a) Defined Gene Sets** |  | **(b) Model Performance** |  |
| Set | Genes | Condition | PR-AUC (%) |
|---|---|---|---|
| A | *panBCD*, *bioABCDFH* | Baseline | 96.1 ± 2.6 |
| B | Set A + *kdsC*, *mdh* | A1 | 96.7 ± 2.1 |
| C | *kdsC*, *mdh* | B1 | 98.1 ± 1.3 |
|  |  | C1 | 97.1 ± 2.1 |
|  |  | A2 | 96.6 ± 2.6 |
|  |  | A2 + C1 | 97.7 ± 1.5 |

#### 3.1.6 Experimental growth assays resolve dataset discrepancies and confirm gene essentiality

To empirically resolve discrepancies between our model predictions and the iML1515 empirical dataset, we performed growth assay experiments on Keio collection single-gene knockout mutants cultured in glucose M9 minimal medium (results in Figure 3). Detailed experimental protocols, pre-conditioning regimes, and kinetic growth parameters estimated using the `fitderiv` package [48] are provided in Appendix B. For Δ*bioF* and Δ*bioH*, direct inoculation from rich Lysogeny Broth (LB) into glucose M9 minimal medium permitted initial proliferation at wild-type levels, sustained by the intracellular pooling and carryover of cofactors and trace metabolic intermediates (Figure 3B). However, a single pre-conditioning passage in minimal medium significantly attenuated growth, and a second consecutive back-dilution into fresh, unsupplemented minimal medium resulted in complete cessation of biomass accumulation (Figure 3), confirming their essentiality in minimal medium. Similarly, upon direct transfer from LB to M9, Δ*panB* supported only marginal initial growth before arresting entirely once residual cofactor pools were exhausted, establishing *panB* as essential. Conversely, Δ*hemF*, Δ*kdsC*, and Δ*mdh* exhibited wild-type-like growth kinetics (Figure 3). Deletion mutants for *ubiH* and *ubiE* displayed compromised fitness, characterized by extended lag phases, reduced maximum optical densities (OD_600_), and lower specific growth rates (Figure 3A), yet remained classified as non-essential because their growth rates stayed above the essentiality threshold (> 50% of the wild-type BW25113 rate).

**Fig. 3:**
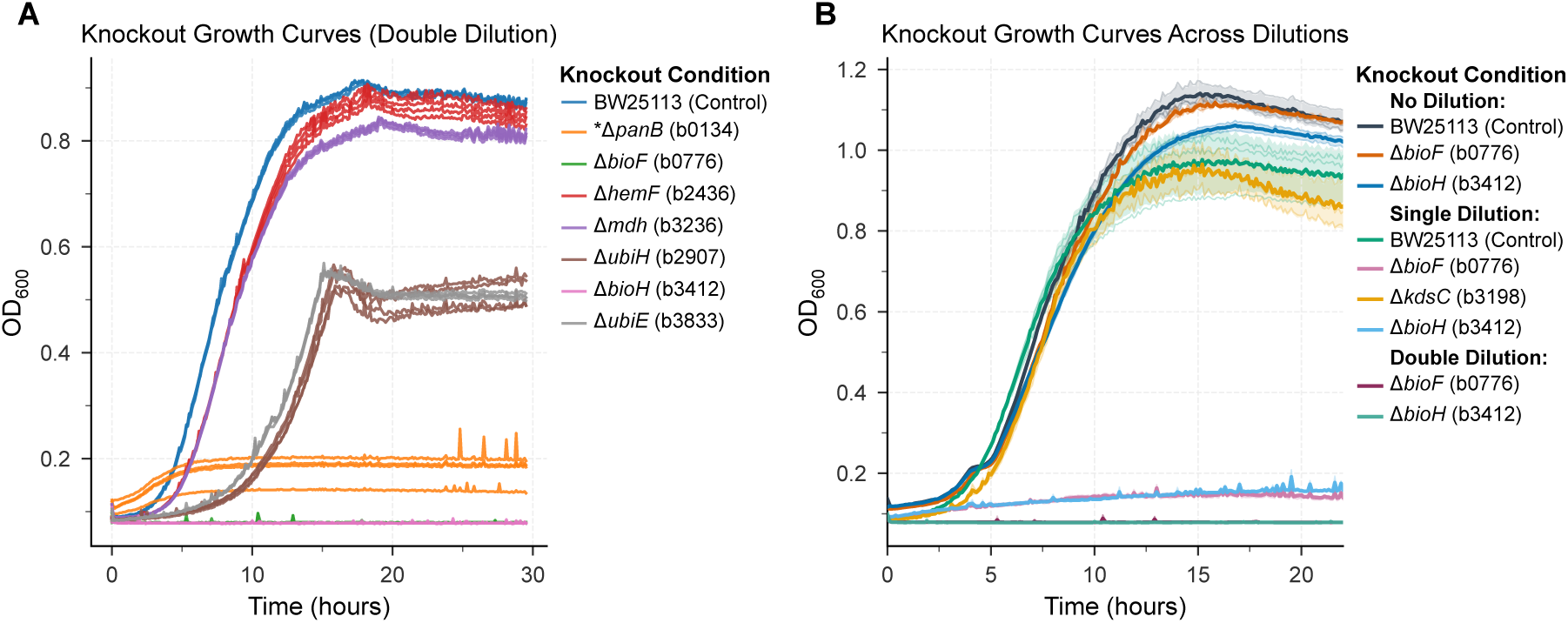
Careful experiments on growth kinetics are required to identify essentiality, avoiding confounding effects from nutritional carryover and pre-conditioning. (**A**), Growth curves of Keio collection knockout strains in M9 glucose minimal medium following double back-dilution. (*): Due to severe growth arrest upon exhaustion of LB-derived cofactors, the auxotroph Δ*panB* was evaluated via direct inoculation from rich medium. (**B**), Pre-conditioning-dependent growth profiles for *kdsC* (b3198), *bioF* (b0776), and *bioH* (b3412) compared to the parental wild-type control (*E. coli* BW25113). While the *kdsC* mutant phenocopies the wild type, *bioF* and *bioH* display robust proliferation following direct inoculation, severely attenuated growth after a single acclimation cycle, and complete growth arrest after secondary back-dilution, demonstrating how residual nutrient carryover masks conditional essentiality. Curves and shaded areas represent the mean ± s.d. of three to four biological replicates.

In summary, the observed growth phenotypes decisively confirmed our model predictions across three disputed genes (Δ*panB*, Δ*bioF*, and Δ*bioH*; with the exception of HyperGenie for *bioF*), demonstrating complete growth arrest in unsupplemented minimal medium and proving that the iML1515 empirical dataset misannotated all three loci as non-essential. We verified the target gene deletions in Δ*panB* and Δ*bioH* by diagnostic colony PCR (Appendix B.2, Figure B3). Together, these findings demonstrate how discrepancies identified by topology-aware machine learning can directly guide the targeted re-curation of reference metabolic reconstructions.

#### 3.1.7 Model performance and generalizability under varying nutrient conditions

Figure 4A demonstrates the ability of HyperGenie to accurately predict gene essentiality across diverse nutrient environments. To assess performance, we compared our method to a no-skill classifier baseline (black dots). Given the inherent imbalance in the dataset, with approximately three times as many essential genes as non-essential genes, this baseline would simply predict every gene to be essential. Our method consistently achieves a substantial improvement in PR-AUC (greater than 12%) compared to this baseline across most of the tested carbon sources.

**Fig. 4:**
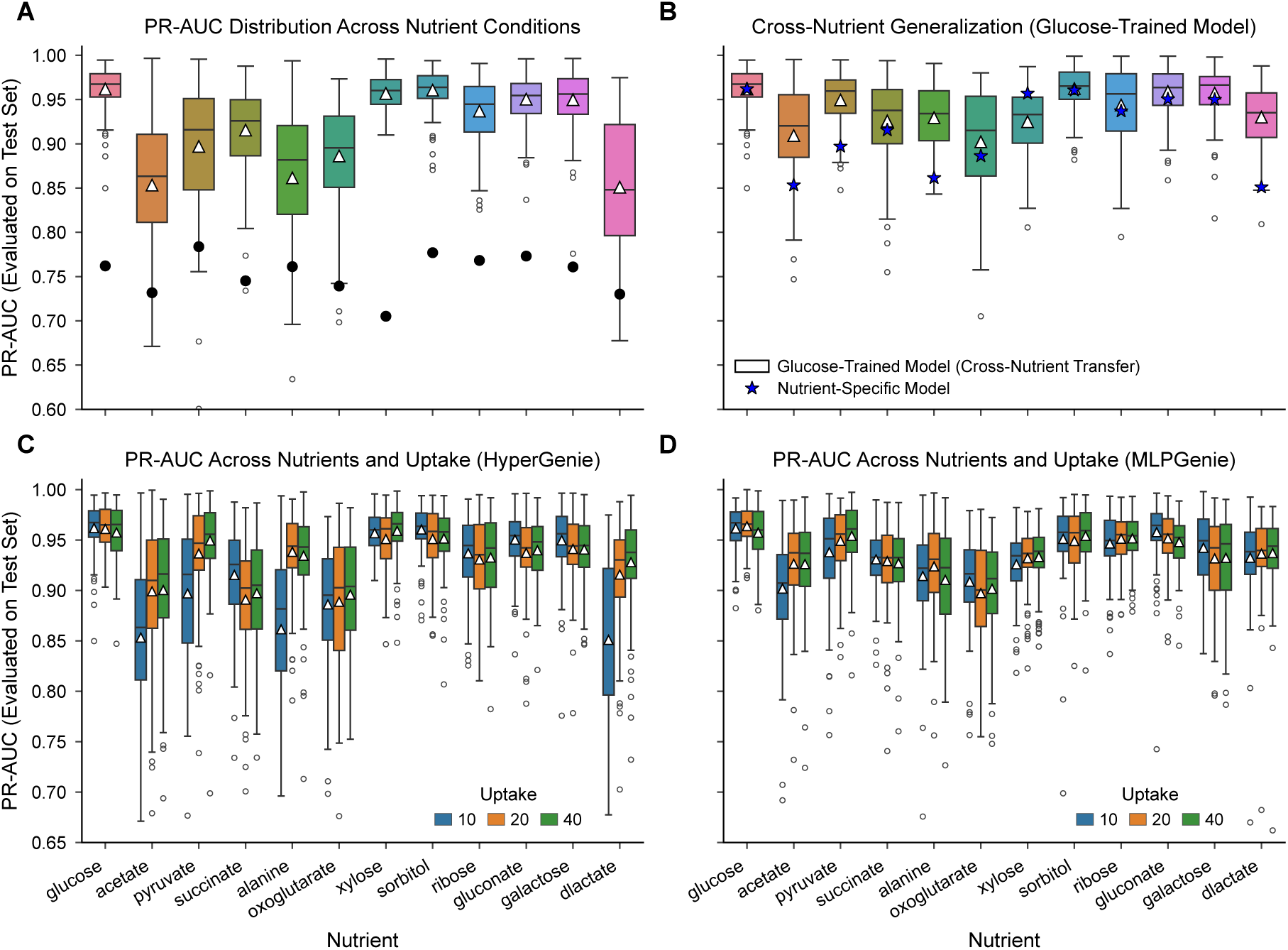
HyperGenie accurately predicts *E. coli* gene essentiality across diverse carbon sources and uptake rates. **(A)** Within-condition predictive performance of HyperGenie for *E. coli* grown aerobically on different sole carbon sources. Boxplots show precision-recall area under the curve (PR-AUC) distributions across 100 runs of 5-fold cross-validation; triangles indicate mean PR-AUC, and black dots denote the baseline PR-AUC without a classifier (demonstrating >15% improvement across most conditions). **(B)** Cross-condition generalization of a glucose-trained HyperGenie model evaluated on alternative carbon sources. Star markers indicate the mean PR-AUC of models trained natively on each target substrate. The glucose-trained model generalizes well to low-molecular-weight substrates but underperforms on xylose. **(C, D)** Sensitivity of predictive performance to carbon uptake rates, evaluated at default (10 mmol gDW^−1^ h^−1^) and elevated (20 and 40 mmol gDW^−1^ h^−1^) bounds across 50 independent runs. **(C)** HyperGenie achieves substantial performance gains on select carbon sources at higher uptake rates, whereas **(D)** MLPGenie exhibits minimal sensitivity to uptake rate, with only acetate showing notable improvement (2.4%).

By utilizing flux features from genome-scale metabolic modelling, HyperGenie and MLPGenie can generalise to any nutrient condition for which experimental data are available. Building on this flexibility, we next investigated whether a model trained exclusively on a single carbon source (glucose) could generalize to predict essentiality in alternative nutrient environments. Such cross-condition predictive capability would significantly expand the model’s utility by circumventing the need for exhaustive, condition-specific experimental datasets.

Figure 4B compares the PR-AUC values obtained from a HyperGenie model trained solely on glucose against HyperGenie models trained natively on the respective target nutrients. HyperGenie exhibits robust cross-condition generalization, with performance on most nutrients statistically indistinguishable from the target-trained baselines. Notably, predictive performance on 2- and 3-carbon sources (Section 3.1.8) is superior when the model is trained on glucose, whereas performance on xylose is significantly diminished.

While overall cross-condition generalization is robust, the core metabolic network remains largely conserved across conditions. Therefore, we next assessed the model’s ability to accurately resolve condition-specific gene essentiality. For this analysis, we used HyperGenie Norm due to its marginally more robust performance across a few of the carbon sources (Section 3.1.8). Table 6 reveals a structural bias in the glucose-trained model: it yields low predictive accuracy for genes that are dispensable in the target nutrient but essential in glucose, yet performs well for genes essential in the target nutrient but dispensable in glucose. This pattern is likely driven, in part, by the model’s general propensity to over-predict gene essentiality.

**Table 6:**
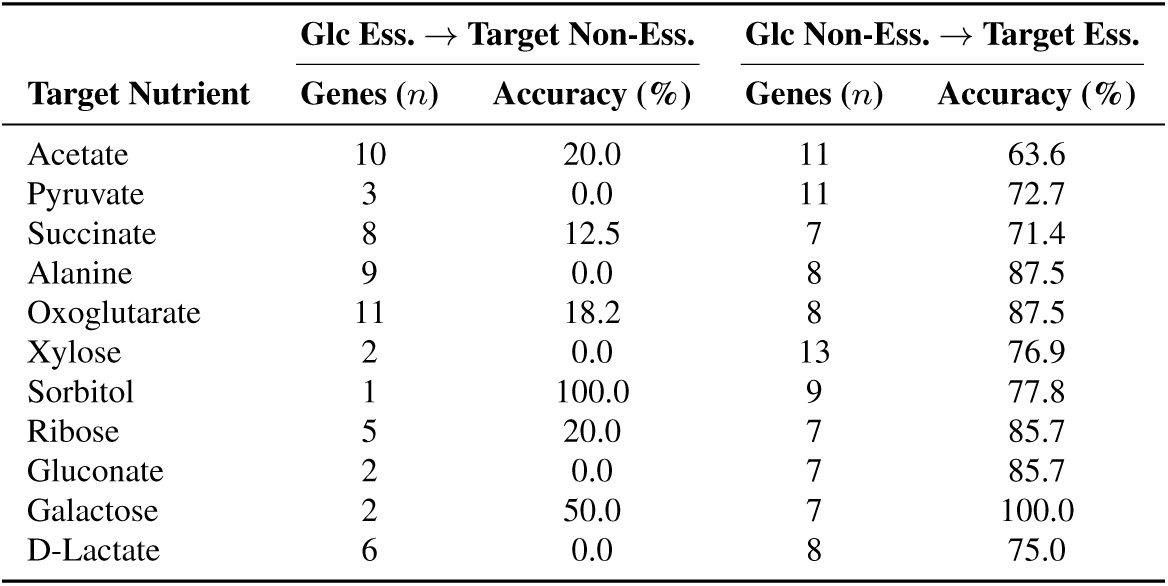
Predictive accuracy of a HyperGenie Norm model trained exclusively on glucose and evaluated on genes exhibiting differential essentiality across 11 alternative carbon sources. Genes are stratified into two categories: those essential in glucose but dispensable in the target environment (left), and those dispensable in glucose but essential in the target environment (right). The model demonstrates high accuracy for genes transitioning to essential, but diminished accuracy for those transitioning to non-essential, reflecting a general tendency to over-predict gene essentiality. A comprehensive list of the specific correctly predicted genes is available in Table D3.

| Target Nutrient | **Glc Ess. → Target Non-Ess.** |  | **Glc Non-Ess. → Target Ess.** |  |
|---|---|---|---|---|
| **Target Nutrient** | **Genes (*n*)** | **Accuracy (%)** | **Genes (*n*)** | **Accuracy (%)** |
| Acetate | 10 | 20.0 | 11 | 63.6 |
| Pyruvate | 3 | 0.0 | 11 | 72.7 |
| Succinate | 8 | 12.5 | 7 | 71.4 |
| Alanine | 9 | 0.0 | 8 | 87.5 |
| Oxoglutarate | 11 | 18.2 | 8 | 87.5 |
| Xylose | 2 | 0.0 | 13 | 76.9 |
| Sorbitol | 1 | 100.0 | 9 | 77.8 |
| Ribose | 5 | 20.0 | 7 | 85.7 |
| Gluconate | 2 | 0.0 | 7 | 85.7 |
| Galactose | 2 | 50.0 | 7 | 100.0 |
| D-Lactate | 6 | 0.0 | 8 | 75.0 |

According to the iML1515 database, 21 genes with a one-to-one or one-to-many relationship to the reactions they catalyse exhibit differential essentiality between glucose and acetate. Seven of these genes are associated with reactions that have zero pFBA-predicted flux in either condition. Although these genes would normally be excluded from our pipeline (Section 2.1), we retained them for this specific analysis. Of the 14 remaining genes that meet our standard inclusion criteria, four shift from non-essential in glucose to essential in acetate, while ten are essential in glucose but dispensable in acetate.

Crucially, the model correctly identified several genes whose essentiality is highly specific to a single carbon source, as identified by the iML1515 database. These include acnB (b0118) in acetate [49], galE (b0759) in galactose [50], xylB (b3564) in xylose [51], and rpsK (b3752) in ribose [52]. Furthermore, it successfully detected genes that are essential across multiple carbon conditions, such as ubiE (b3833) and ubiH (b2907). The full list of correctly predicted differentially essential genes, focusing on those that are dispensable in glucose but essential in the alternative carbon sources, is provided in Table D3. By accurately resolving both condition-specific and broadly essential genes, this approach establishes itself as a powerful tool for discovering novel antibiotic targets across a diverse range of metabolic environments.

#### 3.1.8 Sensitivity to increased nutrient uptake rates

Initial evaluations (Figure 4A) revealed a performance disparity across the different nutrient conditions, with specific carbon sources exhibiting a PR-AUC reduction of over 5% relative to the top-performing substrates. To determine whether these predictive deficits were driven by overly restrictive flux constraints, we increased the carbon nutrient uptake rates.

Figures 4C and 4D show the PR-AUC performance of HyperGenie and MLPGenie, respectively, across simulated uptake rates of 10 (the default), 20 and 40 mmol gDW^−1^ h^−1^. Increasing the uptake rate from 10 to 20 mmol gDW^−1^ h^−1^ substantially rescued HyperGenie’s performance in the previously underperforming environments. Specifically, we observed substantial improvements in PR-AUC when simulating growth on alanine (+7.7%), D-lactate (+6.5%), acetate (+4.6%), and pyruvate (+4.0%). HyperGenie Norm demonstrated a qualitatively similar, albeit attenuated, response (Figure D3). By contrast, while MLPGenie exhibited parallel directional trends, its performance deficit was marginal; across all tested conditions, only acetate yielded an absolute PR-AUC increase exceeding 1.5% (+2.4%). Notably, the predictive performance for the TCA cycle intermediates oxoglutarate and succinate remained largely invariant and persistently low, despite increased uptake limits across both models. Figure D3 in the appendix shows the results in greater detail.

To understand what drives these predictions, we mapped the shift in PR-AUC (comparing the default uptake rate of 10 mmol gDW^−1^ h^−1^ with the elevated rate of 20 mmol gDW^−1^ h^−1^) against substrate molar mass and carbon chain length, and categorized the results by the metabolic entry point of each nutrient (Figure 5). This analysis revealed a pronounced predictive deficit for low-molecular-weight substrates, specifically those comprising two or three carbon atoms. The metabolic routing of these short-chain carbon sources diverges fundamentally from that of five- and six-carbon carbohydrates (e.g., glucose, xylose, sorbitol, ribose, gluconate, and galactose). Growth on these smaller substrates relies heavily on gluconeogenesis to synthesize essential sugar-phosphate biomass precursors. In the case of acetate, assimilation additionally necessitates flux through the glyoxylate shunt to bypass oxidative decarboxylation steps and replenish TCA intermediates. These differences result in more restrictive metabolic fluxes across the whole network, and this appears intrinsically more difficult for HyperGenie to accurately learn.

**Fig. 5:**
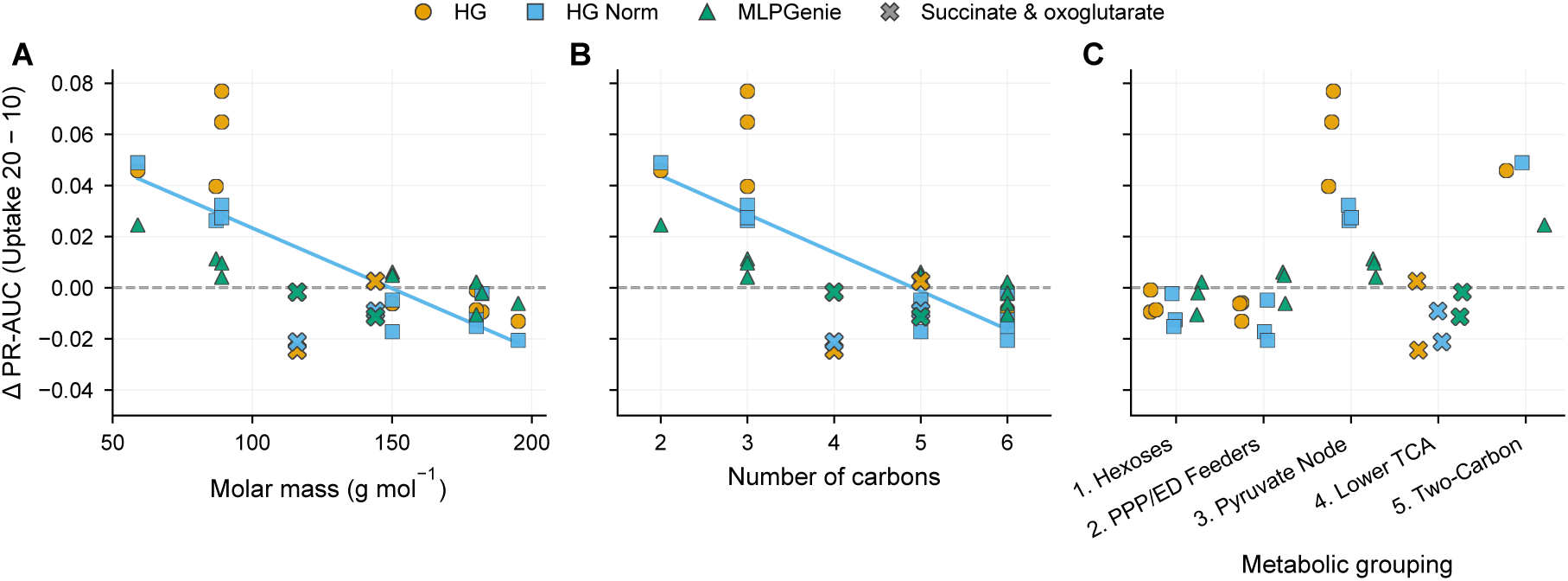
Low-molecular-weight carbon sources exhibit the greatest predictive improvement at elevated uptake rates. Scatter plots depict the change in performance (ΔPR-AUC) when the simulated carbon uptake rate is increased from the default 10 mmol gDW^−1^ h^−1^ to 20 mmol gDW^−1^ h^−1^. Data are shown for HyperGenie (HG), HyperGenie with normalized edge features (HG norm), and MLPGenie, differentiated by marker color and shape. The invariant TCA cycle intermediates, oxoglutarate and succinate, are distinctly denoted by cross markers. **A:** ΔPR-AUC as a function of substrate molar mass. Smaller substrates demonstrate substantially larger performance gains. A reference linear regression line is fitted to the HG norm data, explicitly excluding the outlier TCA cycle intermediates. **B:** ΔPR-AUC versus the number of carbon atoms per substrate, reflecting a trend consistent with molar mass. **C:** Substrates categorized by metabolic group. Short-chain carbon nutrients requiring gluconeogenic flux (e.g., acetate and pyruvate) display the most substantial performance enhancements with increased uptake, an effect particularly pronounced in the HyperGenie models. This sensitivity to uptake constraints arises because short-chain carbon sources (C2–C3) rely heavily on gluconeogenesis and specialized bypass pathways (e.g., the glyoxylate shunt), which enforce tightly constrained network-wide flux distributions that are intrinsically harder for HyperGenie to capture.

#### 3.1.9 Model comparison and performance metrics

Figure 6 summarizes the predictive accuracy of the three models (HyperGenie, HyperGenie Norm, and MLP-Genie) across 12 nutrients at uptake rates of 10, 20, and 40 mmol gDW^−1^ h^−1^. Here, each model was trained and evaluated strictly within individual nutrient conditions without cross-condition generalization. With the exception of the lowest uptake rate (10 mmol gDW^−1^ h^−1^) for the two- and three-carbon nutrients discussed in Section 3.1.8 (acetate, pyruvate, alanine, and D-lactate), both HyperGenie variants consistently outperform MLPGenie. This demonstrates that modelling higher-order interactions is critical for more accurately capturing the underlying biological complexity. Specifically, the hypergraph architecture improved predictive accuracy by at least 2% across most nutrients, with gains exceeding 4% for acetate, pyruvate, oxoglutarate, and ribose, and reaching over 10% for xylose.

**Fig. 6:**
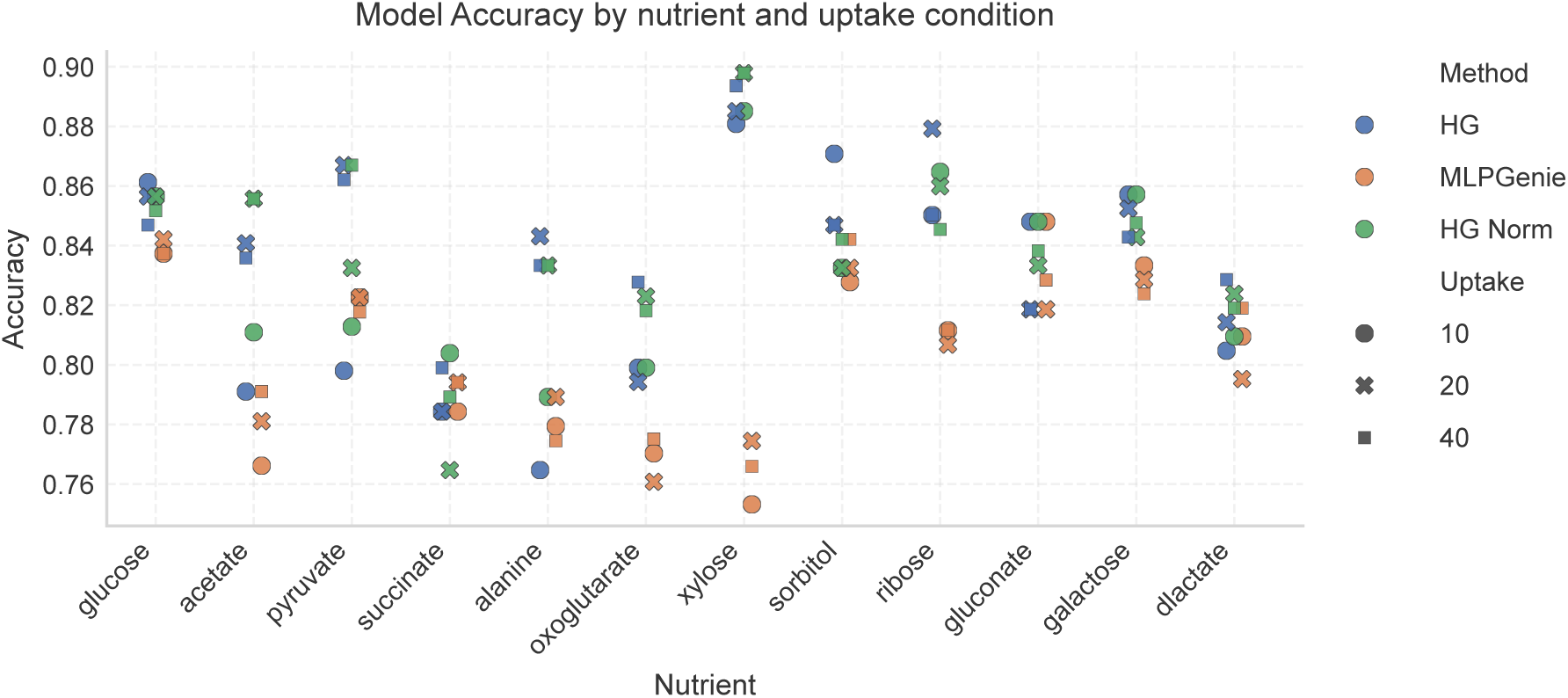
The HyperGenie models have the best accuracy performance across diverse carbon environments. Accuracy metrics for gene essentiality predictions in *E. coli* simulated under aerobic growth with 12 distinct sole carbon sources. Models (HyperGenie (HG), HyperGenie Norm, and MLPGenie) were evaluated at uptake rates of 10 (default), 20, and 40 mmol gDW^−1^ h^−1^. Across most conditions, both HyperGenie variants significantly outperform MLPGenie, yielding absolute accuracy improvements of ≥2% for the majority of nutrients and exceeding 10% for xylose. The sole exception occurs at the lowest uptake rate (10 mmol gDW^−1^ h^−1^) for specific three-carbon nutrients (pyruvate, alanine, and D-lactate). Overall, these results demonstrate that modelling higher-order interactions is critical for accurately capturing the underlying biological complexity.

To provide a more comprehensive evaluation, Table 7 compares the models using additional key metrics (precision, recall, F1 score, and total predicted essential genes) under specific nutrient conditions (e.g., acetate and pyruvate). Overall, the HyperGenie models outperformed MLPGenie. The only exception was observed during simulated growth on pyruvate at an uptake rate of 10 mmol/gDW/h, where MLPGenie performed better than or equal to the HyperGenie models.

**Table 7:** Predictive metrics for acetate and pyruvate across varying carbon uptake rates. Performance comparison of the HyperGenie model, HyperGenie Norm, and MLPGenie. *E. coli* growth was simulated on acetate and pyruvate at substrate uptake rates of 10, 20, and 40 mmol gDW^−1^ h^−1^. The HyperGenie architectures generally outperform MLPGenie across most metrics and uptake conditions, with the exception of pyruvate at 10 mmol gDW^−1^ h^−1^, where MLPGenie performs equally well or better. The total number of true essential genes in the dataset is 150 for Acetate and 159 for Pyruvate.

| Uptake | Model | Precision | Accuracy | Recall | F1 Score | Essential Genes Found |
|---|---|---|---|---|---|---|
| **Acetate** |  |  |  |  |  |  |
| 10 | HyperGenie | 82.1 | 79.1 | 92.0 | 86.8 | 138 |
|  | HyperGenie Norm | 84.6 | 81.1 | 91.3 | 87.8 | 137 |
|  | MLPGenie | 82.8 | 76.6 | 86.7 | 84.7 | 130 |
| 20 | HyperGenie | 86.4 | 84.1 | 93.3 | 89.7 | 140 |
|  | HyperGenie Norm | 89.5 | 85.6 | 91.3 | 90.4 | 137 |
|  | MLPGenie | 84.4 | 78.1 | 86.7 | 85.5 | 130 |
| 40 | HyperGenie | 86.8 | 83.6 | 92.0 | 89.3 | 138 |
|  | HyperGenie Norm | 88.1 | 85.6 | 93.3 | 90.6 | 140 |
|  | MLPGenie | 86.0 | 79.1 | 86.0 | 86.0 | 129 |
| **Pyruvate** |  |  |  |  |  |  |
| 10 | HyperGenie | 83.9 | 79.8 | 91.8 | 87.7 | 146 |
|  | HyperGenie Norm | 85.8 | 81.3 | 91.2 | 88.4 | 145 |
|  | MLPGenie | 87.3 | 82.3 | 90.6 | 88.9 | 144 |
| 20 | HyperGenie | 90.2 | 86.7 | 93.1 | 91.6 | 148 |
|  | HyperGenie Norm | 87.9 | 83.3 | 91.2 | 89.5 | 145 |
|  | MLPGenie | 88.2 | 82.3 | 89.3 | 88.8 | 142 |
| 40 | HyperGenie | 89.2 | 86.2 | 93.7 | 91.4 | 149 |
|  | HyperGenie Norm | 89.8 | 86.7 | 93.7 | 91.7 | 149 |
|  | MLPGenie | 87.7 | 81.8 | 89.3 | 88.5 | 142 |

#### 3.1.10 Explainability

In this section, all reaction and metabolite identifiers, including their shorthand notations, follow the conventions of the *i*ML1515 genome-scale metabolic model from the BiGG Models database [44]. All models were trained and evaluated on *E. coli* grown aerobically in glucose-supplemented M9 minimal medium, except for the data in Figure 8A, where cross-condition evaluations on acetate were performed using glucose-trained model weights coupled with acetate-specific flux features.

##### MLPGenie model

To determine the metabolic network properties that most strongly dictate gene essentiality, we analysed the MLPGenie predictions using SHapley Additive exPlanations (SHAP) [43] via the expected gradients method [53] (detailed methodologies are provided in Appendices C and C.2). The SHAP summary analysis (Figures 7A and 7B) reveals that the neural network prioritises network rigidity over parsimonious flux assumptions. The minimum guaranteed flux required to maintain 99% of the optimal growth (FVA Min) and the standard optimal flux (FBA) emerged as the most impactful predictors, yet they exert opposing directional influences. Specifically, the model learned to associate zero or negative FVA Min values with non-essentiality, while identifying the highest FVA Min values as strict indicators of essential reactions. From a theoretical standpoint, any FVA Min value strictly greater than zero should denote an essential reaction, assuming a flawlessly constructed metabolic network. The model likely learned complex, non-linear relationships between these features; for instance, identifying strong metabolic bottlenecks when FVA Max and FVA Min are closely aligned, or recognising redundant pathways when high FBA values coincide with low FVA Min. Furthermore, higher parsimonious FBA (pFBA) values exhibited a slight pull towards essentiality, suggesting that reactions maintaining high flux even under strict whole-network flux minimisation constraints may possess an underlying biological criticality.

**Fig. 7:**
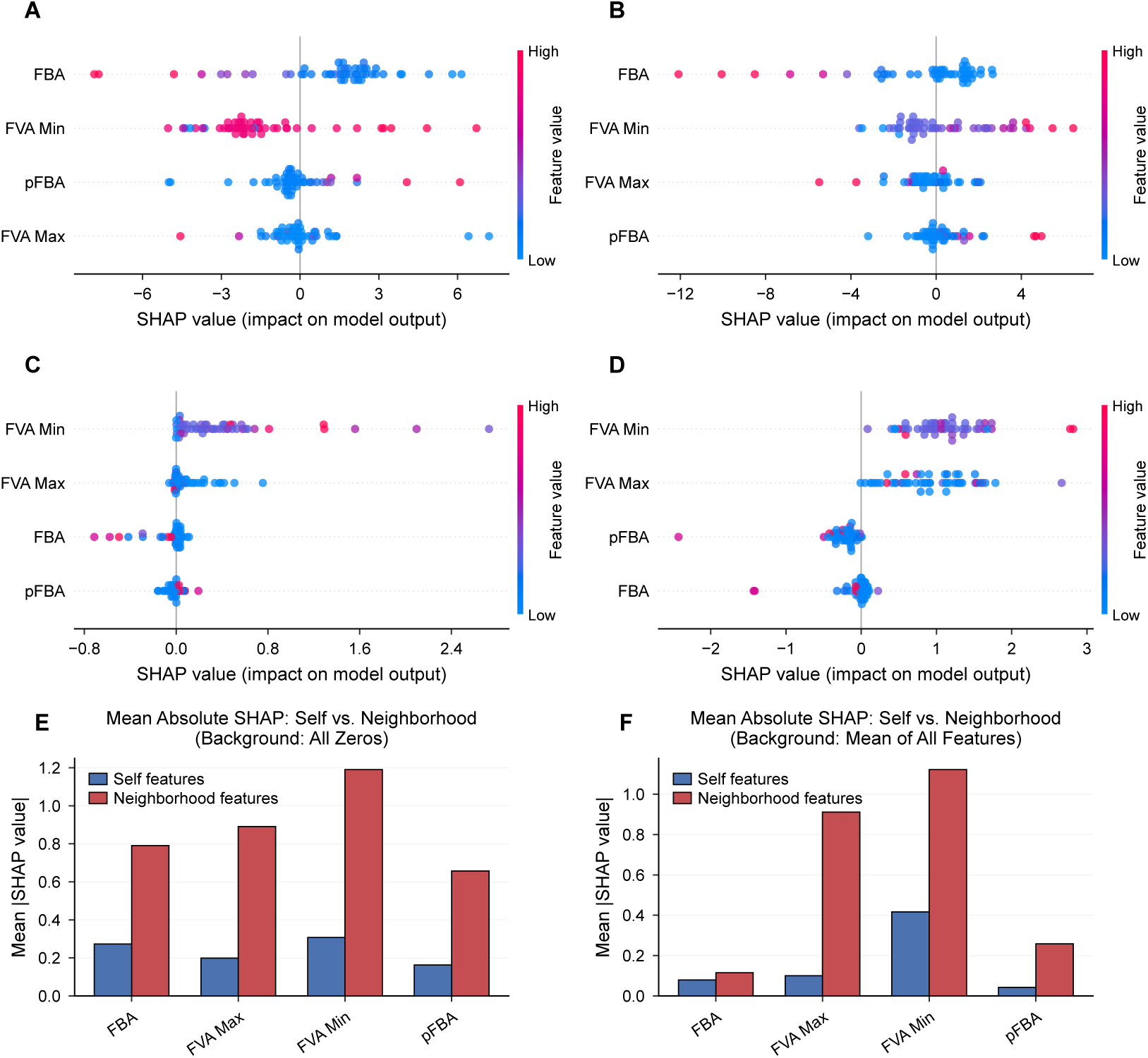
SHAP feature attribution analysis shows that both models prioritise metabolic network rigidity over parsimonious flux assumptions, while HyperGenie heavily weights neighbourhood context over intrinsic reaction features to predict gene essentiality. All models were trained and evaluated on *E. coli* grown aerobically in glucose-supplemented M9 minimal medium. (**A, B**) Global feature impact for the MLP-Genie model across two distinct cross-validation folds, achieving (**A**) 94% and (B) 98% PR-AUC. Minimum guaranteed flux (FVA Min) and standard optimal flux (FBA) consistently emerge as the most impactful predictors. While high FVA Min strongly indicates essential reactions, high FBA indicates non-essential reactions. (**C, D**) HyperGenie SHAP attributions detailing the influence of intrinsic versus network features against a mean background reference. (**C**) Intrinsic (self) features: The impact of a target reaction’s own four features. Higher FVA Min strongly shifts predictions toward essentiality, while high FBA fluxes drive predictions toward non-essentiality. (**D**) Aggregated network (neighborhood) features: The influence of the wider metabolic environment, calculated for each target reaction as the signed sum of SHAP attributions across all other network reactions. While local directional trends are conserved, the network context exhibits greater variance, with FVA Max surpassing FBA in importance. Baseline reference values: FBA = 0.0799, FVA Max = 0.9796, FVA Min = −0.2782, pFBA = 0.0800. (**E, F**) Sensitivity of global feature importance to the SHAP background reference across *n* = 50 test samples (target reactions), comparing (**E**) an all-zero baseline against (**F**) the mean training feature baseline. Bars represent the mean of the absolute SHAP values across the 50 test samples for the intrinsic and aggregated neighborhood attributions defined in (**C, D**). Across both baselines, FVA bounds (FVA Min and FVA Max) dominate the attribution signal, and the cumulative neighborhood contribution substantially outweighs that of the intrinsic features, confirming that HyperGenie relies primarily on wider network context to predict essentiality.

##### HyperGraph explainability

Figures 7C and 7D present the SHAP summary plots detailing the feature attributions derived from a reaction’s own intrinsic features and the aggregated features of all other reactions in the network, respectively. SHAP values were calculated using the expected gradients method. The baseline reference consisted of the mean flux values across the entire graph, excluding the test set (FBA = 0.0799, FVA Max = 0.9796, FVA Min = −0.2782, pFBA = 0.0800).

Unlike MLPGenie, the HyperGenie model incorporates topological context through hypergraph message passing. This allows it to shift its predictive focus primarily toward flux capacity boundaries (FVA Min and FVA Max). This transition makes biological sense: while FBA only offers a single optimal snapshot, FVA bounds reveal the flexibility and redundancy of the metabolic network. Directionally, because the background reference for FVA Min is negative (−0.2782), all FVA Min values appear to shift the model’s predictions toward essentiality, with higher minimum fluxes exerting a stronger positive impact. Conversely, high FBA values tend to drive predictions towards non-essentiality. Interestingly, pFBA demonstrates a diverging effect between intrinsic and neighbourhood features. While a high intrinsic pFBA value pulls the model toward predicting essentiality, a high aggregated pFBA from the surrounding neighbourhood pushes it toward non-essentiality. Perhaps this points to a dynamic learned by the model: a highly efficient, high-flux reaction is critical on its own, but if it is situated within a highly active and redundant metabolic neighbourhood, the broader local network is robust enough to withstand its deletion.

Because SHAP values are inherently relative to the reference data, these findings, derived by generating gradients against a mean background, must be interpreted exclusively within the context of this chosen baseline strategy. Figures 7E and 7F present the mean absolute SHAP values for each feature, comparing the attributions derived from a target reaction’s own features against the aggregated SHAP values of features from all other reactions in the network. Unlike MLPGenie, HyperGenie makes predictions by incorporating this broader network context alongside a reaction’s internal features. To evaluate how the baseline reference impacts these interpretations, we compared two distinct approaches. Figure 7E illustrates the results when applying a background strategy of all zeros, whereas Figure 7F displays the outcomes when using the mean of all training features as the background. As these plots demonstrate, shifting the background baseline significantly alters the resulting global feature attributions.

To identify the reactions driving the model’s overall decision-making, we calculated an aggregated global SHAP magnitude. For each test prediction, we first determined the local impact of every background reaction by averaging the absolute SHAP values across its four features. We then established global reaction importance by averaging these local magnitudes across all out-of-fold test predictions from our 5-fold cross-validation (*n* = 254 for glucose and *n* = 270 for acetate). The larger reaction set in the acetate condition reflects the inclusion of reactions carrying non-zero pFBA flux that remain inactive (zero flux) under glucose conditions. This approach highlights the reactions that most strongly dictate essentiality predictions across the entire metabolic network (Figure 8A).

**Fig. 8:**
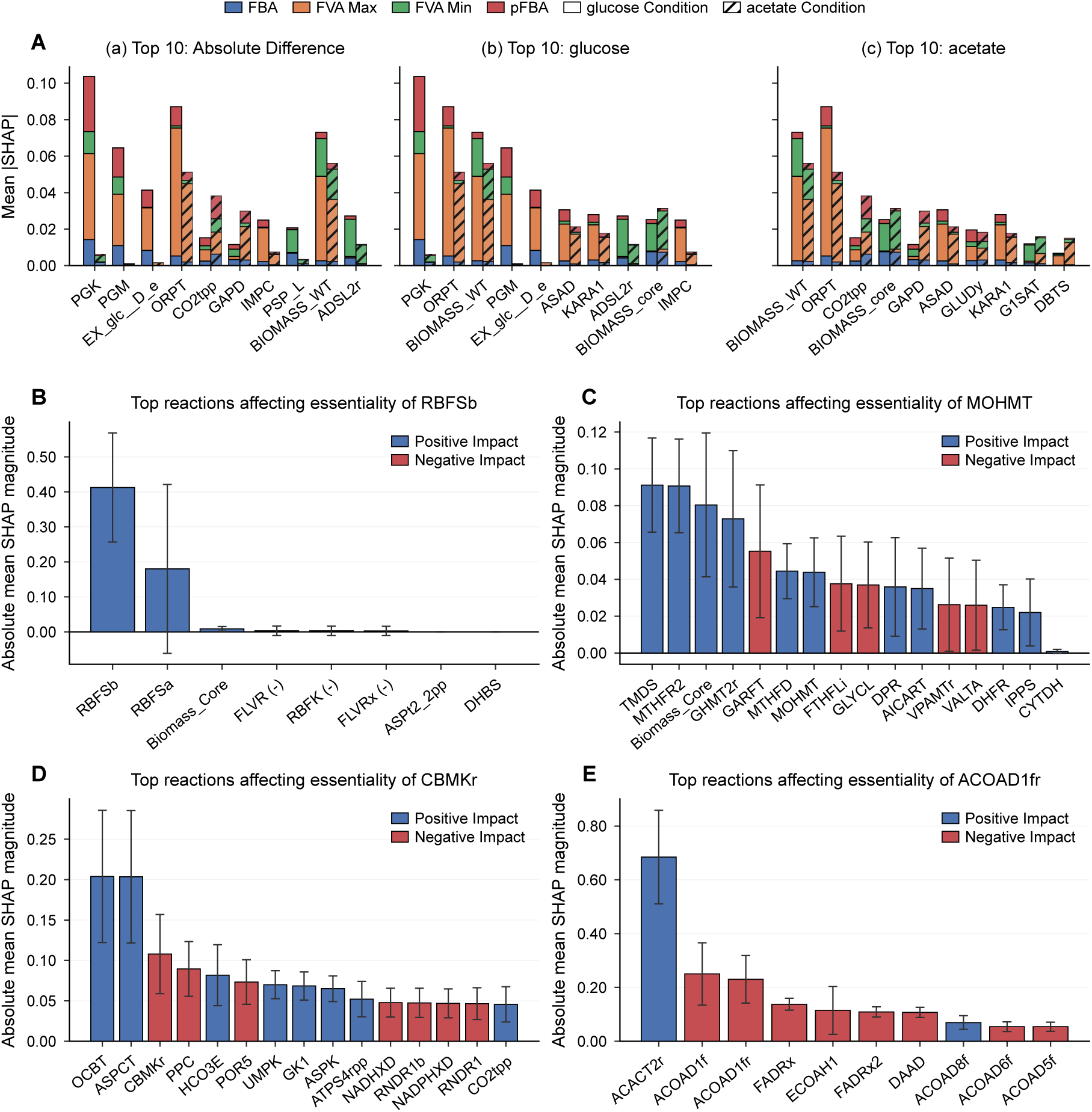
Global and local SHAP feature attributions confirm that HyperGenie learns biologically consistent representations of global flux rewiring and local reaction dependencies. Models were trained on *E. coli* in glucose M9 minimal medium; acetate results in (**A**) represent cross-condition evaluations using acetate flux features, while (**B–E**) show predictions under glucose conditions. (A) Global reaction importance across nutrients. (a) Top ten reactions exhibiting the largest shift in mean absolute SHAP values between glucose and acetate. (b, c) Top ten most globally influential reactions evaluated under (b) glucose and (c) acetate. Feature attributions capture key metabolic adaptations, highlighting the structural transition from downward glycolysis in glucose to upward gluconeogenesis in acetate (e.g., PGK, PGM, GAPD), the carbon-conserving glyoxylate shunt (CO_2_tpp), and downstream biosynthetic demands such as nucleotide (ORPT, IMPC, ADSL2r) and amino acid (PSPL) pathways. (**B–E)** Local network drivers of individual predictions. Top metabolic drivers for individual queried reactions (error bars: s.d. across 10 random seeds). (**B**) Essential RBFSb relies on its upstream precursor RBFSa. (**C**) MOHMT essentiality is governed by one-carbon folate pool dynamics. (**D**) Non-essential ACOAD1fr dependencies reflect fatty acid metabolism and functional competition for the shared oxidized FAD pool. (E) Non-essential CBMKr depends on carbamoyl phosphate sinks, CO_2_ cycling, and energy balance.

Figure 8A highlights the ten reactions with the greatest difference in SHAP magnitude between glucose and acetate, while panels (b) and (c) display the top ten most important reactions for each respective carbon source. The results are highly consistent with established biological principles, demonstrating that HyperGenie captures an accurate representation of metabolic network dynamics. Notably, the model successfully detects the systemic shift in central carbon directionality. Reactions such as PGK, PGM, EX glc D e, and GAPD^2^ are central to this divergence. Glucose metabolism drives high, downward fluxes through glycolysis to generate abundant energy and pyruvate. In contrast, acetate enters the metabolic network downstream as acetyl-CoA; to synthesize the essential hexose and pentose sugars required for biomass, the cell must drive carbon upward via gluconeogenesis. This reversal transforms PGK, PGM, and GAPD into tightly constrained bottlenecks during acetate metabolism, whereas carbon flows through them with much higher redundancy in glucose. Furthermore, the reaction CO2tpp (CO_2_ transporter via diffusion (periplasm)) emerges as a primary discriminator reflecting carbon scarcity. The standard tricarboxylic acid (TCA) cycle loses two carbon atoms as CO_2_ for every molecule of acetyl-CoA oxidized. Because acetate provides exactly two carbons, utilizing the full TCA cycle would result in zero net carbon assimilation. Consequently, acetate-grown cells route a portion of metabolic flux through the glyoxylate shunt to bypass these oxidative decarboxylation steps, balancing energy generation with the retention of essential carbon for biomass, a strict conservation mechanism that is entirely bypassed in a carbon- and energy-rich glucose environment.

Beyond central carbon routing, the model also captures the heightened sensitivity of biosynthetic pathways, particularly amino acid and nucleotide synthesis, under carbon limitation. The differential importance of the phosphoserine phosphatase reaction (PSP L) serves as a prime example. L-serine (ser__L c) is derived from 3-phosphoglycerate (3pg c). During glucose metabolism, 3pg c is a highly abundant glycolytic intermediate; however, during growth on acetate, it must be synthesized via energy-intensive gluconeogenesis. This upstream scarcity makes the downstream consumption of 3pg c highly sensitive, elevating PSP L to a structurally critical node. Similarly, the high rankings of purine and pyrimidine biosynthesis reactions (ORPT, IMPC, ADSL2r)^3^ highlight the metabolic strain of nucleotide production on suboptimal carbon sources. Nucleotide synthesis requires a constant supply of ribose-5-phosphate (r5p c) alongside immense, direct ATP investments for base polymerization. While glucose directly and efficiently feeds the oxidative pentose phosphate pathway (PPP), growth on acetate forces the network to route flux upward via energy-intensive gluconeogenesis to generate upper-glycolytic precursors, namely fructose-6-phosphate (f6p c) and glyceraldehyde-3-phosphate (g3p c), before they can be transformed into r5p c. This extended, thermodynamically demanding route, coupled with the lower overall energy yield of acetate oxidation, severely constrains nucleotide production. Consequently, reactions like ORPT, IMPC, and ADSL2r become highly sensitive metabolic vulnerabilities within the acetate essentiality network.

##### Local explanations

A key capability of our framework is the extraction of local explanations, identifying the specific metabolic reactions that drive the model’s essentiality prediction for any single queried reaction. To achieve this, we computed local feature attributions using the Kernel SHAP estimator (60,000 iterations; detailed methodology in Appendix C.1). We applied a binary mask to scale the feature vector of each individual reaction. The baseline (background) distribution was defined by a zero-mask across all reactions, effectively simulating complete feature absence (i.e., all hyperedge features set to zero). Conversely, test samples were evaluated using a mask of ones to preserve the original edge features, representing the intact HyperGenie model.

To demonstrate this capability, we profiled four test cases (reaction shorthand notations follow [44]). Two of these are strictly essential for *E. coli* survival in glucose M9 minimal media: RBFSb (riboflavin synthase) and MOHMT (3-methyl-2-oxobutanoate hydroxymethyltransferase). RBFSb, encoded by *ribC* (b1662), catalyzes the terminal step of *de novo* riboflavin biosynthesis; lacking a human homolog, it serves as an attractive antimicrobial target. MOHMT, encoded by *panB* (b0134), is indispensable for pantothenate biosynthesis. We contrasted these with two reactions that are non-essential under identical minimal media conditions: CBMKr (carbamate kinase) and ACOAD1fr (acyl-CoA dehydrogenase), the latter of which functions in membrane lipid metabolism (Figures 8B–E).

Analysis of RBFSb essentiality across these ten replicates revealed a tightly coupled structural dependency (Figure 8B). In seven of the ten replicates, the hyperedge features for RBFSb and its immediate upstream precursor, RBFSa (lumazine synthase), exhibited the highest and most comparable SHAP values, indicating that the model heavily prioritizes this linked biosynthetic module. In the remaining three replicates, the model assigned the majority of the predictive weight directly to the RBFSb hyperedge, with RBFSa yielding near-zero or negative SHAP values; this variance accounts for the high standard deviation observed in Figure 8B. Minor SHAP contributions were also attributed to the biomass objective function and downstream flavin-processing reactions (FLVR, RBFK, and FLVRx)^4^ which collectively act as metabolic sinks driving riboflavin consumption. Ultimately, the robust predictive coupling between RBFSb and RBFSa accurately reflects their strict stoichiometric codependence as RBFSa is the sole generator of 6,7-dimethyl-8-(1-D-ribityl)lumazine, the only requisite substrate of RBFSb. Furthermore, the condensation of two lumazine molecules by RBFSb yields riboflavin alongside a secondary byproduct that can be recycled directly back into the RBFSa reaction. For the MOHMT reaction (Eq. (4)), the highest-ranking network features by absolute mean SHAP magnitude reveal a strict structural dependency on the cellular one-carbon folate pool (Figure 8C). Specifically, the most predictive reactions are topologically coupled to the cycling of 5,10-methylenetetrahydrofolate (mlthf) and tetrahydrofolate (thf). The model assigns the largest SHAP magnitudes to major metabolic sinks that actively consume mlthf, namely TMDS, MTHFR2, MTHFD, and the biomass core reaction (BIOMASS Ec iML1515 core 75p37M). Concurrently, substantial predictive weight is allocated to GHMT2r (Eq. (5)) and GLYCL, which utilize thf to replenish the mlthf pool via distinct pathways. Furthermore, the model captures broader folate dynamics by highlighting distal thf-generating reactions (e.g., GARFT (Eq. (6)), AICART, and DHFR) alongside competing thf-consuming sinks (FTHFLi) as critical modulators of MOHMT essentiality.^5^

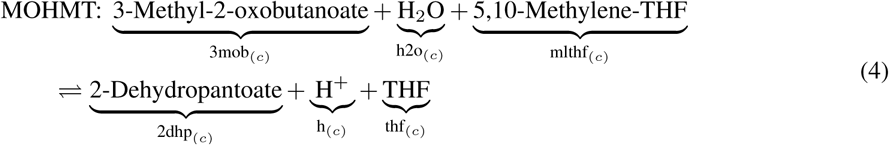

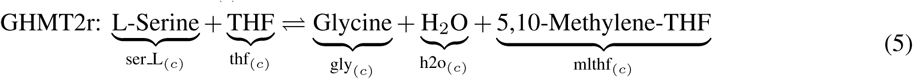

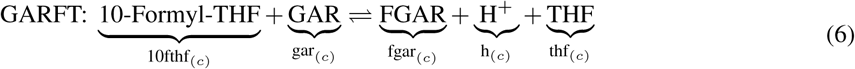

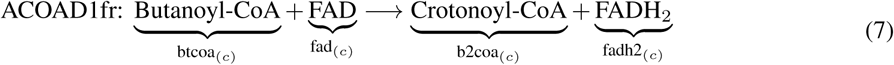

Analysis of the CBMKr (carbamate kinase) reaction reveals a primary predictive dependence on the cellular demand for carbamoyl phosphate (Figure 8D). The model assigns the highest SHAP magnitudes to aspartate carbamoyltransferase (ASPCT) and ornithine carbamoyltransferase (OCBT), which serve as the principal downstream sinks consuming carbamoyl phosphate for pyrimidine and arginine biosynthesis, respectively. Beyond these direct topological successors, the network captures upstream substrate dependencies by allocating significant predictive weight to the carbon dioxide sub-network. Specifically, reactions governing CO_2_ consumption, production, and periplasmic transport (PPC, HCO3E, POR5, and CO2tpp)^6^ act as critical distal modulators. Furthermore, the model highlights key nodes governing the global ATP/ADP balance, capturing both major energy-consuming kinases (UMPK, GK1, and ASPK) and the primary ATP synthase (ATPS4rpp)^7^. Together, these attributions demonstrate that CBMKr essentiality is tightly coupled not only to direct biosynthetic sinks, but also to global CO_2_ cycling and the cellular energy charge.

Analysis of the acyl-CoA dehydrogenase reaction (ACOAD1fr, Eq. (7)) reveals network dependencies that align with the topology of fatty acid metabolism (Figure 8E). The dominant predictive contribution arises from Acetyl-CoA C-acyltransferase:butanoyl-CoA (ACACT2r). Because ACACT2r consumes butanoyl-CoA (btcoa), it functions as a critical metabolic sink that relies directly on the forward flux of ACOAD1fr; accordingly, the model assigns it a strongly positive SHAP value. Conversely, substantial negative predictive weights are allocated to both ACOAD1f, the direct thermodynamic reverse of the target reaction, and ECOAH1 (3-hydroxyacyl-CoA dehydratase:3-hydroxybutanoyl-CoA), which synthesizes the requisite crotonoyl-CoA substrate upstream. Furthermore, the model highlights a broader suite of peripheral oxidoreductases (including FADRx, FADRx2, DAAD, ACOAD6f, and ACOAD5f)^8^ that uniformly exert negative SHAP contributions. By competitively reducing the shared cytosolic pool of oxidized FAD to FADH_2_, these secondary reactions act in functional opposition to ACOAD1fr.

## 4 Conclusion

The environmental and nutritional dependency of gene essentiality poses a major bottleneck for experimental validation, underscoring the need for robust computational prediction methods. While genome-scale metabolic models (GEMs) provide a powerful predictive framework, conventional constraint-based methods like flux balance analysis (FBA) assume that mutant strains continue to optimize for maximum growth. However, this assumption may not always be valid, as gene knockouts can disrupt cellular processes and lead to suboptimal or alternative growth strategies. Prior machine learning approaches, such as random forests and graph neural networks utilizing Mass Flow Graphs (MFGs), have attempted to bypass this limitation by training on wild-type fluxes. However, these methods fundamentally compromise network topology: random forests often discard spatial structural context entirely, while MFGs are restricted to pairwise connections, failing to capture the multi-reactant, multi-product dependencies inherent to biochemistry. To resolve this, we introduced HyperGenie: a hypergraph neural network that explicitly models these higher-order systemic dependencies by mapping the *E. coli* metabolic network into a hypergraph topology. Crucially, because the entire metabolic network is structurally embedded within this representation, HyperGenie provides an interpretable framework, enabling us to identify the specific metabolic reactions driving the model’s essentiality predictions.

Operating directly on wild-type flux distributions, HyperGenie bypasses the restrictive maximal-growth assumptions applied to knockout strains, outperforming previous objective-dependent graph methods by 13% in PR-AUC while achieving accuracy, precision, and F1-scores that match or exceed the standard FBA baseline. HyperGenie also achieves performance parity with Flux Cone Learning (FCL), the current state of the art. While FCL eliminates the need for a predefined objective function via flux sampling, a crucial advantage for modelling eukaryotic systems, the sampling process inherently requires additional computational resources. Moreover, because FCL relies on a random forest classifier, it is significantly less amenable to mechanistic interpretation. HyperGenie, therefore, offers a direct alternative that preserves topological inductive biases without requiring sampling overhead.

To isolate the value of hypergraph message passing, we engineered MLPGenie, a multilayer perceptron baseline, demonstrating that while global constraint-based features (FVA bounds) enable an MLP to achieve comparable PR-AUC, HyperGenie consistently provides superior classification accuracy (>2% gain in glucose and >4% across alternative carbon sources). Crucially, both MLPGenie and HyperGenie demonstrate exceptional sample efficiency, maintaining robust predictive performance (PR-AUC >90%) using only 20% and 40% of the training data, respectively. Beyond benchmark metrics, HyperGenie exhibits strong out-of-distribution generalization. Zero-shot predictions, training exclusively on glucose data and predicting across diverse alternative nutrients, yielded performance equivalent to models trained directly in their native environments. Evaluating the model across varying uptake rates revealed robust performance across most conditions, with a notable exception for two- and three-carbon nutrients; here, performance degraded at standard uptake rates (10 mmol/gDW/h) but was mostly recovered at higher rates (20 or 40 mmol/gDW/h). A deeper analysis of conditionally essential genes highlighted that HyperGenie can accurately predict genes that gain essentiality in novel environments, though it struggles to predict the loss of essentiality. Furthermore, leveraging systematic discrepancies between our model predictions and the iML1515 empirical dataset exposed gene essentiality misannotations in that dataset within the pantothenate (vitamin B_5_) and biotin (vitamin B_7_) pathways. Guided by these discrepancies, we used targeted serial-dilution growth assays to experimentally resolve three disputed genes (Δ*panB*, Δ*bioF*, and Δ*bioH*), demonstrating how model-guided analysis can correct errors in reference datasets and expose experimental growth artifacts.

Finally, to bridge computational predictions with biological mechanism, multi-scale SHAP analyses established the interpretability of HyperGenie at both global and local resolutions. Globally, feature attributions confirmed that the architecture relies primarily on FVA flux bounds to predict essentiality. In addition, despite being trained exclusively on aerobic glucose M9 minimal medium, the model generalized to acetate M9 conditions to autonomously capture key mechanistic adaptations: the directional reversal of central carbon flux between glycolysis and gluconeogenesis, carbon conservation via the glyoxylate shunt, and emergent biosynthetic vulnerabilities under two-carbon substrate limitation. Locally, Kernel SHAP demonstrated that gene essentiality is governed by multi-tier metabolic coupling, linking the prioritization of immediate precursor supply, competitive downstream sinks of key intermediates, and distal modulators regulating shared cofactor pools.

Despite these advantages, our framework possesses specific limitations that outline clear directions for future research. First, the current iteration of HyperGenie relies on an objective function to derive wild-type fluxes. However, the hypergraph architecture itself is agnostic to the underlying flux generation method; future implementations can be adapted to use flux sampling distributions, enabling its application to complex eukaryotic models. Second, the current model predicts essentiality at the reaction level and subsequently maps these predictions back to specific genes via gene-protein-reaction (GPR) rules. This structural workflow currently limits its application to genes with straightforward one-to-one or one-to-many relationships with metabolic reactions.

## Supplementary information

Supplementary information accompanying this paper is available as an appendix in a separate document.

## Data and code availability

The complete dataset and implementation code for this work will be openly accessible in the Zenodo repository under https://zenodo.org/records/23955982. All computational workflows were executed on a local workstation running Linux Mint 21.3, equipped with an Intel Core i7-13700 CPU (16 cores, 24 threads), 16 GB DDR4 RAM, and an NVIDIA GeForce RTX 3060 GPU (12 GB VRAM, CUDA 12.4). Under this configuration, completing 10 independent iterations of 5-fold cross-validation (50 total model training and evaluation runs) required approximately 9 minutes in total (approximately 10.8 seconds per individual run).

## Acknowledgements

P.I. was supported by the Engineering and Physical Sciences Research Council (EPSRC) Centre for Doctoral Training in Sensor Technologies for a Healthy and Sustainable Future [grant number EP/S023046/1]. P.I. also gratefully acknowledges additional financial support from the Cambridge Philosophical Society via a three-month research studentship, and from Robinson College, Cambridge, via a two-month research studentship.

P.I. thanks Leonardo Mancini for technical guidance and advice regarding bacterial culture protocols, growth assays, and kinetic curve analyses. The authors also thank Abhirup Mookherjee for guidance and assistance with the PCR verification experiments. We are grateful to Teresa Saez and the Sainsbury Laboratory (University of Cambridge) for providing the *E. coli* Keio collection knockout strains used in this work.

C.A. was supported by UKRI BBSRC (grant BB/Y01278X/1) and Horizon Europe (grant 101182278). Gemini (Google) was used to refine grammar, clarity, and the visual formatting of selected tables and figures. All final content was critically reviewed, edited, validated, and approved by the authors.

## Author contributions

P.I.: Conceptualization, Data curation, Formal analysis, Investigation, Methodology, Software, Validation, Visualization, Writing – original draft, Writing – review & editing. I.D.: Methodology, Validation, Writing – review & editing. S.V.: Methodology, Validation, Writing – review & editing. P.C.: Conceptualization, Methodology, Supervision, Writing – review & editing. P.L.: Conceptualization, Methodology, Supervision, Writing – review & editing. C.A.: Conceptualization, Methodology, Project administration, Supervision, Writing – review & editing.

## Footnotes

1 Some metabolites in the metabolic network, such as the proton (H+), are highly connected and participate in hundreds of biochemical reactions. The feature generation method described above could therefore result in extremely high counts for these nodes. To mitigate the potential for these high counts to disproportionately influence training, we set an arbitrary maximum count of one million for any feature that has a higher count.

2 PGK: phosphoglycerate kinase; PGM: phosphoglycerate mutase; EX_glc_D_e: D-glucose exchange; GAPD: glyceraldehyde-3-phosphate dehydrogenase.

3 ORPT: orotate phosphoribosyltransferase; IMPC: inosine monophosphate cyclohydrolase; ADSL2r: adenylosuccinate lyase.

4 FLVR: flavin reductase; RBFK: riboflavin kinase; FLVRx: flavin reductase (NAD).

5 TMDS: thymidylate synthase; MTHFR2: 5,10-methylenetetrahydrofolate reductase (NADH); MTHFD: methylenetetrahydrofolate dehydrogenase (NADP); BIOMASS_Ec_iML1515_core_75p37M: core biomass objective function; GHMT2r: glycine hydroxymethyltransferase; GLYCL: glycine cleavage system; GARFT: phosphoribosylglycinamide formyltransferase; AICART: phosphoribosylaminoimidazolecarboxamide formyltransferase; DHFR: dihydrofolate reductase; FTHFLi: formate-tetrahydrofolate ligase.

6 PPC: phosphoenolpyruvate carboxylase; HCO3E: HCO3^−^ equilibration reaction; POR5: pyruvate synthase.

7 UMPK: uridylate kinase; GK1: guanylate kinase; ASPK: aspartate kinase; ATPS4rpp: ATP synthase (four protons per ATP, periplasm to cytoplasm).

8 FADRx: flavin adenine dinucleotide reductase; FADRx2: flavin adenine dinucleotide reductase (NADH/NADPH); DAAD: D-amino acid dehydrogenase; ACOAD6f: acyl-CoA dehydrogenase (tetradecanoyl-CoA); ACOAD5f: acyl-CoA dehydrogenase (dodecanoyl-CoA).

## References

[1] Fang, G., Rocha, E., Danchin, A.: How essential are nonessential genes? Molecular biology and evolution 22(11), 2147–2156 (2005)

[2] Rancati, G., Moffat, J., Typas, A., Pavelka, N.: Emerging and evolving concepts in gene essentiality. Nature Reviews Genetics 19(1), 34–49 (2018)

[3] Aromolaran, O., Aromolaran, D., Isewon, I., Oyelade, J.: Machine learning approach to gene essentiality prediction: a review. Briefings in bioinformatics 22(5), 128 (2021)

[4] Li, X., Li, W., Zeng, M., Zheng, R., Li, M.: Network-based methods for predicting essential genes or proteins: a survey. Briefings in bioinformatics 21(2), 566–583 (2020)

[5] Schapke, J., Tavares, A., Recamonde-Mendoza, M.: EPGAT: gene essentiality prediction with graph attention networks. IEEE/ACM Transactions on Computational Biology and Bioinformatics 19(3), 1615–1626 (2021)

[6] Zhang, X., Xiao, W., Xiao, W.: DeepHE: Accurately predicting human essential genes based on deep learning. PLOS Computational Biology 16(9), 1008229 (2020)

[7] Hasan, M.A., Lonardi, S.: DeeplyEssential: a deep neural network for predicting essential genes in microbes. BMC bioinformatics 21, 1–19 (2020)

[8] Occhipinti, A., Hamadi, Y., Kugler, H., Wintersteiger, C.M., Yordanov, B., Angione, C.: Discovering essential multiple gene effects through large scale optimization: an application to human cancer metabolism. IEEE/ACM transactions on computational biology and bioinformatics 18(6), 2339–2352 (2020)

[9] Nandi, S., Ganguli, P., Sarkar, R.R.: Essential gene prediction using limited gene essentiality information–An integrative semi-supervised machine learning strategy. PLoS One 15(11), 0242943 (2020)

[10] Hwang, Y., Lin, C., Chang, J., Mori, H., Juan, H., Huang, H.: Predicting essential genes based on network and sequence analysis. Molecular BioSystems 5(12), 1672–1678 (2009)

[11] Liu, X., He, T., Guo, Z., Ren, M., Luo, Y.: Predicting essential genes of 41 prokaryotes by a semi-supervised method. Analytical biochemistry 609, 113919 (2020)

[12] Campos, T.L., Korhonen, P.K., Sternberg, P.W., Gasser, R.B., Young, N.D.: Predicting gene essentiality in Caenorhabditis elegans by feature engineering and machine-learning. Computational and Structural Biotechnology Journal 18, 1093–1102 (2020)

[13] Ye, Y., Zhou, R., Li, L., Yuan, H., Xia, J., Li, Y., Zeng, Z., Zhang, X.: CNN4Essential: a convolutional neural network model for predicting bacterial gene essentiality based on multi-feature fusion. BMC Genomics (2026)

[14] Kang, B., Fan, R., Cui, C., Cui, Q.: Comprehensive prediction and analysis of human protein essentiality based on a pretrained large language model. Nature Computational Science 5(3), 196–206 (2025)

[15] Mansouri-Rad, S., Narimani, Z., Razzaghi, P., Hosseinkhan, N.: EssentialGIN: a new approach for gene essentiality prediction based on graph isomorphism neural networks. arXiv preprint arXiv:2606.07700 (2026)

[16] Kim, B.K., Gu, C., Farh, M.E., Ryu, J.Y.: Integrating Genome-Scale Metabolic Modeling with Machine Learning Improves Gene Essentiality Prediction in Triple-Negative Breast Cancer. International Journal of Molecular Sciences 27(11), 5059 (2026)

[17] Rahimi, S., Bonner, S., Afzal, A., Milo, M., Morrissey, E., Petsalaki, E.: Biological Network Organization, Not Generic Graph Topology, Drives Graph-Based Gene Essentiality Prediction. bioRxiv, 2026–06 (2026)

[18] O’Brien, E.J., Monk, J.M., Palsson, B.O.: Using genome-scale models to predict biological capabilities. Cell 161(5), 971–987 (2015)

[19] Monk, J.M., Lloyd, C.J., Brunk, E., Mih, N., Sastry, A., King, Z., Takeuchi, R., Nomura, W., Zhang, Z., Mori, H., et al.: iML1515, a knowledgebase that computes Escherichia coli traits. Nature biotechnology 35(10), 904–908 (2017)

[20] Joyce, A.R., Palsson, B.Ø.: Predicting gene essentiality using genome-scale in silico models. Microbial Gene Essentiality: Protocols and Bioinformatics, 433–457 (2008)

[21] Orth, J.D., Thiele, I., Palsson, B.Ø.: What is flux balance analysis? Nature biotechnology 28(3), 245–248 (2010)

[22] Mahadevan, R., Schilling, C.H.: The effects of alternate optimal solutions in constraint-based genome-scale metabolic models. Metabolic engineering 5(4), 264–276 (2003)

[23] Segre, D., Vitkup, D., Church, G.M.: Analysis of optimality in natural and perturbed metabolic networks. Proceedings of the national academy of sciences 99(23), 15112–15117 (2002)

[24] Hasibi, R., Michoel, T., Oyarzún, D.A.: Integration of graph neural networks and genome-scale metabolic models for predicting gene essentiality. npj Systems Biology and Applications 10(1), 24 (2024)

[25] Freischem, L.J., Oyarzún, D.A.: A Machine Learning Approach for Predicting Essentiality of Metabolic genes. In: Synthetic Biology: Methods and Protocols, pp. 345–369. Springer, New York, NY (2023)

[26] Beguerisse-Dıaz, M., Bosque, G., Oyarzún, D., Pico, J., Barahona, M.: Flux-dependent graphs for metabolic networks. NPJ systems biology and applications 4(1), 32 (2018)

[27] Schuetz, R., Kuepfer, L., Sauer, U.: Systematic evaluation of objective functions for predicting intracellular fluxes in Escherichia coli. Molecular systems biology 3, 119 (2007)

[28] Feist, A.M., Palsson, B.O.: The biomass objective function. Current opinion in microbiology 13(3), 344–349 (2010)

[29] Schinn, S., Morrison, C., Wei, W., Zhang, L., Lewis, N.E.: Systematic evaluation of parameters for genome-scale metabolic models of cultured mammalian cells. Metabolic engineering 66, 21–30 (2021)

[30] Merzbacher, C., Mac Aodha, O., Oyarzún, D.A.: Accurate prediction of gene deletion phenotypes with Flux Cone Learning. Nature Communications 16(1), 8492 (2025)

[31] Klamt, S., Haus, U., Theis, F.: Hypergraphs and cellular networks. PLoS computational biology 5(5), 1000385 (2009)

[32] Gallo, G., Longo, G., Pallottino, S., Nguyen, S.: Directed hypergraphs and applications. Discrete applied mathematics 42(2-3), 177–201 (1993)

[33] Battiston, F., Cencetti, G., Iacopini, I., Latora, V., Lucas, M., Patania, A., Young, J., Petri, G.: Networks beyond pairwise interactions: Structure and dynamics. Physics reports 874, 1–92 (2020)

[34] Traversa, P., Arruda, G., Vazquez, A., Moreno, Y.: Robustness and complexity of directed and weighted metabolic hypergraphs. Entropy 25(11), 1537 (2023)

[35] Feng, Y., You, H., Zhang, Z., Ji, R., Gao, Y.: Hypergraph neural networks. In: Proceedings of the AAAI Conference on Artificial Intelligence, vol. 33, pp. 3558–3565 (2019)

[36] Su, G., Wang, H., Zhang, Y., Wilkins, M.R., Canete, P.F., Yu, D., Yang, Y., Zhang, W.: Inferring gene regulatory networks by hypergraph generative model. Cell Reports Methods 5(4) (2025)

[37] Huang, Y., Liang, X., Lin, T., Liu, J.: Multi-HGNN: Multi-modal hypergraph neural networks for predicting missing reactions in metabolic networks. Information Sciences 704, 121960 (2025)

[38] Zhao, Y., Chen, Y., Yu, Y., Liu, X., Du, J., Wen, J., Sun, Q., Wang, R., Chen, C.: A multi-way SMILES-based hypergraph inference network for metabolic model reconstruction. Communications Biology 9(1), 531 (2026)

[39] Chen, S., Chen, T., Xu, Z., Zhang, L., Gao, B., Mao, J.: CytoGem-XAI: A Hypergraph Neural Network Framework for Genome-Scale Metabolic Modeling and Interpretable Analysis. bioRxiv, 2026–06 (2026)

[40] Ioannou, P., Duta, I., Verma, S., Cicuta, P., Lio, P., Angione, C.: Hypergenie: A method for predicting enzymatic gene essentiality using hypergraph neural networks and genome-scale metabolic models. In: ICLR 2025 Workshop on AI for Nucleic Acids

[41] Chien, E., Pan, C., Peng, J., Milenkovic, O.: You are allset: A multiset function framework for hypergraph neural networks. arXiv preprint arXiv:2106.13264 (2021)

[42] Cooper, K., Barahona, M.: Role-based similarity in directed networks. arXiv preprint arXiv:1012.2726 (2010)

[43] Lundberg, S.M., Lee, S.: A unified approach to interpreting model predictions. Advances in neural information processing systems 30 (2017)

[44] King, Z.A., Lu, J., Drager, A., Miller, P., Federowicz, S., Lerman, J.A., Ebrahim, A., Palsson, B.O., Lewis, N.E.: BiGG Models: A platform for integrating, standardizing and sharing genome-scale models. Nucleic acids research 44(D1), 515–522 (2016)

[45] Orth, J.D., Conrad, T.M., Na, J., Lerman, J.A., Nam, H., Feist, A.M., Palsson, B.Ø.: A comprehensive genome-scale reconstruction of Escherichia coli metabolism—2011. Molecular systems biology 7, 535 (2011)

[46] Patrick, W.M., Quandt, E.M., Swartzlander, D.B., Matsumura, I.: Multicopy suppression underpins metabolic evolvability. Molecular biology and evolution 24(12), 2716–2722 (2007)

[47] Sperandeo, P., Pozzi, C., Deho, G., Polissi, A.: Non-essential KDO biosynthesis and new essential cell envelope biogenesis genes in the Escherichia coli yrbG–yhbG locus. Research in microbiology 157(6), 547–558 (2006)

[48] Swain, P.S., Stevenson, K., Leary, A., Montano-Gutierrez, L.F., Clark, I.B., Vogel, J., Pilizota, T.: Inferring time derivatives including cell growth rates using Gaussian processes. Nature communications 7(1), 13766 (2016)

[49] Gruer, M.J., Bradbury, A.J., Guest, J.R.: Construction and properties of aconitase mutants of Escherichia coli. Microbiology 143(6), 1837–1846 (1997)

[50] Lee, S.J., Trostel, A., Le, P., Harinarayanan, R., FitzGerald, P.C., Adhya, S.: Cellular stress created by intermediary metabolite imbalances. Proceedings of the National Academy of Sciences 106(46), 19515–19520 (2009)

[51] Rosenfeld, S.A., Stevis, P.E., Ho, N.W.: Cloning and characterization of the xyl genes from Escherichia coli. Molecular and General Genetics MGG 194(3), 410–415 (1984)

[52] Lopilato, J.E., Garwin, J., Emr, S., Silhavy, T., Beckwith, J.: D-ribose metabolism in Escherichia coli K-12: genetics, regulation, and transport. Journal of bacteriology 158(2), 665–673 (1984)

[53] Erion, G., Janizek, J.D., Sturmfels, P., Lundberg, S.M., Lee, S.: Improving performance of deep learning models with axiomatic attribution priors and expected gradients. Nature machine intelligence 3(7), 620–631 (2021)

