## Appendix for "HyperGenie: A method for predicting enzymatic gene essentiality using hypergraph neural networks and genome-scale metabolic models"

#### Appendix A GEMs, flux balance analysis and flux variability analysis

Genome-scale metabolic models (GEMs), such as *i*ML1515 [1] for *Escherichia coli* K-12 MG1655, provide a comprehensive computational framework capturing the totality of an organism's known biochemical reactions, reconstructed from annotated genome sequences and experimentally validated literature. Mathematically, a GEM is formalized around a stoichiometric matrix $\mathbf{S} \in \mathbb{R}^{m\times n}$, where entries represent the stoichiometric coefficients of $m$ metabolites across $n$ biochemical reactions. Embedded Gene-Protein-Reaction (GPR) associations use Boolean logic to map metabolic genes directly to enzyme complexes and their corresponding catalysed reactions, enabling the simulation of phenotypic behaviours without requiring detailed enzymatic kinetic parameters. Each reaction $j$ is subject to thermodynamic and environmental capacity constraints:

$$v_{j,\mathrm{lb}} \le v_j \le v_{j,\mathrm{ub}}, \quad \forall j \in \{1, \dots, n\} \tag{A1}$$

where $v_{j,\mathrm{lb}}$ and $v_{j,\mathrm{ub}}$ denote the lower and upper flux bounds, respectively. Irreversible reactions enforce $v_{j,\mathrm{lb}} = 0$, whereas exchange flux bounds define nutrient availability in the extracellular medium (e.g., aerobic glucose M9 minimal medium). Cellular proliferation is represented by a biomass objective function, a pseudo-reaction that stoichiometrically accounts for essential macromolecular precursors, including amino acids, nucleotides, lipids, and cofactors, alongside growth-associated and non-growth-associated ATP maintenance requirements.

To predict metabolic phenotypes, Flux Balance Analysis (FBA) [2] employs linear programming to compute an optimal steady-state flux distribution. In this study, the objective is the maximization of the specific growth rate ($\mu$), corresponding to the flux through the biomass pseudo-reaction ($Z = v_{\mathrm{biomass}}$). Under the quasi-steady-state assumption, intracellular metabolite pools remain constant over time ($\mathbf{Sv} = \mathbf{0}$). The linear programming problem is formulated as:

$$
\begin{aligned}
\max_{\mathbf{v}} \quad & Z = \mathbf{c}^T \mathbf{v} \\
\text{subject to} \quad & \mathbf{S}\mathbf{v} = \mathbf{0}, \\
& v_{j,\text{lb}} \leq v_j \leq v_{j,\text{ub}}, \quad \forall j \in \{1, \dots, n\}
\end{aligned} \tag{A2}
$$

where $\mathbf{v} \in \mathbb{R}^n$ represents the vector of metabolic fluxes ($\text{mmol} \cdot \text{gDW}^{-1} \cdot \text{h}^{-1}$) and $\mathbf{c} \in \mathbb{R}^n$ is a vector of linear objective weights, structured here as a standard unit vector selecting $v_{\text{biomass}}$ such that $\mathbf{c}^T\mathbf{v} = v_{\text{biomass}}$.

Because genome-scale networks are inherently underdetermined ($n > m$), the linear programming problem frequently exhibits solution degeneracy, yielding multiple distinct combinations of internal reaction fluxes that achieve the exact same optimal objective value ($Z_{\text{opt}}$). Consequently, standard FBA provides a single representative snapshot of metabolic activity from an entire polyhedral optimal space. To evaluate the full spectrum of allowable flux states, Flux Variability Analysis (FVA) computes the minimum and maximum permissible fluxes through each individual reaction $j$ while preserving a defined fraction of the maximal growth rate:

$$
\begin{aligned}
\min_{\mathbf{v}} / \max_{\mathbf{v}} \quad & v_j \\
\text{subject to} \quad & \mathbf{S}\mathbf{v} = \mathbf{0}, \\
& \mathbf{c}^T\mathbf{v} \geq \gamma Z_{\text{opt}}, \\
& v_{j,\text{lb}} \leq v_j \leq v_{j,\text{ub}}, \quad \forall j \in \{1, \dots, n\}
\end{aligned} \tag{A3}
$$

where $\gamma \in [0, 1]$ designates the fraction of the optimal objective required to be maintained (set to $\gamma = 0.99$ in this study). The resulting intervals $[v_{j,\text{lb}}^{\text{FVA}}, v_{j,\text{ub}}^{\text{FVA}}]$ define the allowable flux range for each reaction consistent with near-optimal growth.

### Appendix B Experimental methods: Bacterial strains and growth assays

*E. coli* single-gene deletion mutants derived from the parent strain BW25113 were sourced from the Keio Collection, kindly provided on Lysogeny Broth (LB) agar plates by The Sainsbury Laboratory. Upon receipt, single colonies were cultured in liquid LB medium and archived as permanent glycerol stocks in 20% (v/v) glycerol stored at -80°C. For routine experimental use, strains were revived by streaking from frozen stocks onto fresh LB agar plates, grown overnight at $37\,^\circ\text{C}$, and maintained as working plates at $4\,^\circ\text{C}$.

To initiate experiments, single colonies were picked from working plates to inoculate liquid LB starter cultures. Keio knockout strains were cultured in LB medium supplemented with kanamycin ($1\times$ working concentration, diluted $1:1000$ from a $1000\times$ stock; $50\,\mu\text{g}\,\text{mL}^{-1}$ final), whereas the parental wild-type control strain (*E. coli* BW25113) was grown in antibiotic-free LB broth due to the absence of the selection cassette. Starter cultures were incubated overnight at $37\,^\circ\text{C}$ with continuous shaking ($200\,\text{rpm}$) and subsequently evaluated under one of three pre-conditioning regimes:

1. **Direct inoculation:** Saturated LB starter cultures were used directly to inoculate experimental growth plates without prior minimal medium acclimation.
2. **Single pre-conditioning (nutritional acclimation):** LB starter cultures were subcultured into $5\,\text{mL}$ of fresh M9 glucose minimal medium in $50\,\text{mL}$ conical tubes (with kanamycin for knockouts; antibiotic-free for BW25113) and incubated overnight ($37\,^\circ\text{C}$, $200\,\text{rpm}$) to adapt cells from rich to minimal media.
3. **Double back-dilution (secondary acclimation):** Following the initial overnight growth in M9 minimal medium (as detailed in Regime 2), cultures were back-diluted into fresh M9 medium and grown overnight for a second consecutive cycle ($37\,^\circ\text{C}$, $200\,\text{rpm}$) to ensure complete nutritional equilibration.

#### B.1 Microplate inoculation and growth kinetics

For all three regimes, kinetic growth profiling was performed in sterile 96-well flat-bottom microplates. Each well was loaded with $200\,\mu\text{L}$ of fresh M9 glucose minimal medium and inoculated with $2\,\mu\text{L}$ of the designated overnight culture ($202\,\mu\text{L}$ total working volume, $\approx 1:100$ dilution). Assays included multiple biological replicates (three or four) per strain alongside wells with only medium inside.

Absorbance at $600\,\text{nm}$ ($OD_{600}$) was monitored continuously using an automated microplate reader maintained at $37\,^\circ\text{C}$ with continuous double-orbital shaking. Measurements were recorded at regular intervals

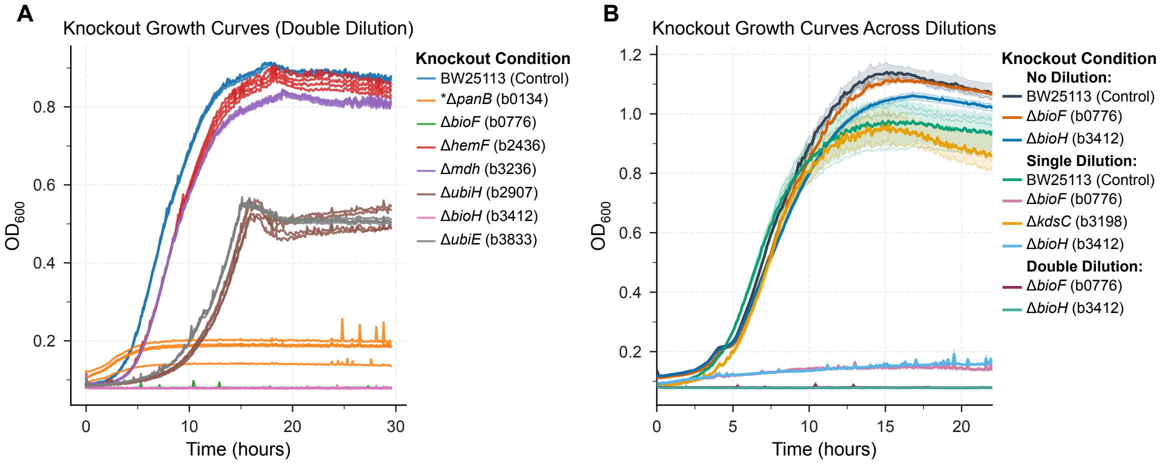


**Fig. B1**: **Careful experiments on growth kinetics are required to identify essentiality, avoiding confounding effects from nutritional carryover and pre-conditioning.** **(A)**, Growth curves of Keio collection knockout strains in M9 glucose minimal medium following double back-dilution. (*): Due to severe growth arrest upon exhaustion of LB-derived cofactors, the auxotroph Δ*panB* was evaluated via direct inoculation from rich medium. **(B)**, Pre-conditioning-dependent growth profiles for *kdsC* (b3198), *bioF* (b0776), and *bioH* (b3412) compared to the parental wild-type control (*E. coli* BW25113). While the *kdsC* mutant phenocopies the wild type, *bioF* and *bioH* display robust proliferation following direct inoculation (Regime 1), severely attenuated growth after a single acclimation cycle, and complete growth arrest after secondary back-dilution, demonstrating how residual nutrient carryover masks conditional essentiality. Curves and shaded areas represent the mean ± s.d. of three to four biological replicates.

(every 6 to 8 mins) over 24 h to 36 h to extract essentiality, maximum specific growth rates ($\mu_{\max}$), and lag phase durations ($\lambda$).

Figure B1A displays representative growth profiles obtained under Regime 3 (double back-dilution), with the exception of Δ*panB*, which was evaluated under Regime 1 (direct inoculation) due to acute growth arrest in minimal medium once residual cofactors from the LB starter culture were exhausted. As shown in Figure B1B, the Δ*kdsC* (b3198) mutant exhibited growth kinetics comparable to the wild-type strain. In contrast, the Δ*bioF* (b0776) and Δ*bioH* (b3412) mutants displayed pronounced, regime-dependent phenotypes across the three pre-conditioning stages: both strains proliferated with near-wild-type kinetics following direct inoculation from LB (Regime 1), exhibited only marginal residual growth after a single round of nutritional acclimation (Regime 2), and arrested completely following the second back-dilution (Regime 3). The corresponding growth kinetic parameters, estimated using the `fitderiv` package [3], are summarized in Table B1, and the derivative growth-rate profiles are shown in Figure B2.

#### B.2 Confirmation of gene knockouts via colony/lysate PCR

##### Template DNA preparation

Bacterial strains (wild-type *E. coli* BW25113, *panB*, and *bioH* knockout mutants) were cultured overnight in LB broth at 37 °C with agitation. Cells were pelleted, washed, and resuspended in sterile nuclease-free water before being lysed at 95 °C for 5 minutes. Following centrifugation to pellet cellular debris, the cleared lysate supernatant was collected as template DNA for downstream PCR.

##### PCR amplification and thermocycling conditions

PCR amplification was conducted to verify target deletions at the *bioH* and *panB* loci. Reactions were prepared in a final volume of 25 $\mu$L, comprising 12.5 $\mu$L of Q5 High-Fidelity 2× Master Mix, 1.25 $\mu$L of each gene-specific forward and reverse primer (10 $\mu$M stock; Table B2), 2.5 $\mu$L of template DNA lysate, and nuclease-free water adjusted to the final volume.

Amplification was performed using the following cycling profile: initial denaturation at 95 °C for 5 minutes; 30 cycles of denaturation at 95 °C for 35 seconds, primer annealing at 63 °C for 30 seconds, and extension at 72 °C for 1 minute; followed by a final extension step at 72 °C for 7 minutes. Samples were subsequently held at 4 °C until analysis.

##### Agarose gel electrophoresis

PCR amplicons were resolved on a 1.0% (w/v) agarose gel prepared in 1× TAE buffer (Thermo Scientific) and

**Table B1**: **Estimated kinetic growth parameters across knockout strains.** Growth parameters were determined using the non-parametric derivative method (`fitderiv`) from $OD_{600}$ time-series data in M9 minimal medium. Data are presented as mean ± s.d. across biological replicates ($N$).

| **Strain / Knockout** | $N$ | **Max Growth Rate** ($h^{-1}$) | **Lag Time** (h) | **Time of $\mu_{max}$** (h) | **Max $OD_{600}$ **** |
|---|---|---|---|---|---|
| **Regime 1: Direct Inoculation (No Acclimation)** | | | | | |
| BW25113 (Control) | 3 | $0.450 \pm 0.002$ | $3.83 \pm 0.07$ | 5.54 | $1.142 \pm 0.024$ |
| Δ*bioF* (b0776) | 3 | $0.455 \pm 0.013$ | $3.66 \pm 0.11$ | 5.82 | $1.116 \pm 0.005$ |
| Δ*bioH* (b3412) | 3 | $0.415 \pm 0.006$ | $3.45 \pm 0.06$ | 5.51 | $1.060 \pm 0.012$ |
| Δ*panB* (b0134)* | 4 | $0.113 \pm 0.020$ | $0.51 \pm 0.22$ | 2.09 | $0.181 \pm 0.027$ |
| **Regime 2: Single Pre-Conditioning (Nutritional Acclimation)** | | | | | |
| BW25113 (Control) | 3 | $0.431 \pm 0.008$ | $2.49 \pm 0.06$ | 4.97 | $0.972 \pm 0.074$ |
| Δ*bioF* (b0776) | 3 | $0.067 \pm 0.003$ | $0.00 \pm 0.00^{\dagger}$ | 0.00 | $0.148 \pm 0.005$ |
| Δ*kdsC* (b3198) | 3 | $0.416 \pm 0.017$ | $2.99 \pm 0.19$ | 5.84 | $0.950 \pm 0.052$ |
| Δ*bioH* (b3412) | 3 | $0.092 \pm 0.006$ | $0.00 \pm 0.00^{\dagger}$ | 0.00 | $0.161 \pm 0.002$ |
| **Regime 3: Double Back-Dilution (Secondary Acclimation)** | | | | | |
| BW25113 (Control) | 4 | $0.382 \pm 0.005$ | $2.59 \pm 0.03$ | 4.89 | $0.906 \pm 0.004$ |
| Δ*bioF* (b0776) | 4 | $0.000 \pm 0.000$ | N/D§ | N/D§ | $0.079 \pm 0.000$ |
| Δ*bioH* (b3412) | 4 | $0.0009 \pm 0.001$ | N/D§ | N/D§ | $0.079 \pm 0.000$ |
| Δ*hemF* (b2436) | 4 | $0.360 \pm 0.005$ | $3.46 \pm 0.07$ | 5.83 | $0.881 \pm 0.013$ |
| Δ*mdh* (b3236) | 4 | $0.372 \pm 0.004$ | $3.93 \pm 0.14$ | 5.58 | $0.833 \pm 0.005$ |
| Δ*ubiH* (b2907) | 4 | $0.239 \pm 0.004$ | $7.85 \pm 0.10$ | 13.91 | $0.535 \pm 0.020$ |
| Δ*ubiE* (b3833) | 4 | $0.242 \pm 0.002$ | $7.30 \pm 0.24$ | 13.33 | $0.552 \pm 0.002$ |

* Δ*panB* was evaluated via direct inoculation (Regime 1) due to rapid cofactor exhaustion leading to total growth arrest upon serial passaging.
† A fitted lag time of zero indicates marginal initial baseline drift/carryover rather than genuine exponential growth.
§ N/D: Not determined. Severe growth inhibition resulted in parameter fitting artifacts.
** Medium $OD_{600}$ included ($0.079 \pm 0.001$)

**Table B2**: Oligonucleotide primers used for diagnostic verification of knockout loci.

| **Target Locus** | **Primer ID** | **Sequence (5′ → 3′)** | **WT Amplicon Size** |
|---|---|---|---|
| *bioH* (771 bp) | KO-bioH-F | `AATCCAAAGCCCTGCATATAG` | 973 bp |
| | KO-bioH-R | `GCATAATCCCGGTACTGTTAG` | |
| *panB* (795 bp) | KO-panB-F | `TCGTCGACCAGCTTCATA` | 1020 bp |
| | KO-panB-R | `CAGTGCTGCTACGGATAAAT` | |

stained with GelRed (Biotium; 1:10,000 dilution). Samples were mixed with orange loading dye, and 10 $\mu$L was loaded per lane. A 100 bp DNA ladder was run in parallel to estimate amplicon sizes. Electrophoresis was carried out at a constant 100 V in 1× TAE buffer until the dye front had migrated across the majority of the gel.

**Knockout verification**

Genotypic validation of the *panB* and *bioH* knockout strains was confirmed via agarose gel electrophoresis (Figure B3). The wild-type control yielded the expected amplicons at 973 bp for *bioH* and 1020 bp for *panB*. In contrast, the Δ*panB* mutant lacked the 1020 bp band and instead displayed a ∼1.3 kb band, consistent with the insertion of the kanamycin resistance cassette ($kan^R$). For the Δ*bioH* mutant, no amplicon was detected, confirming the absence of the target bioH coding sequence.

### Appendix C SHAP (SHapley Additive exPlanations)

Shapley values were originally introduced by Shapley [4] in cooperative game theory as an axiomatic method to fairly distribute a total payout among collaborating players based on their marginal contributions. Lundberg and Lee [5] adapted this framework for machine learning interpretability by framing model prediction as a cooperative game: the input features act as players, and the model prediction $f(x)$ represents the total payout.

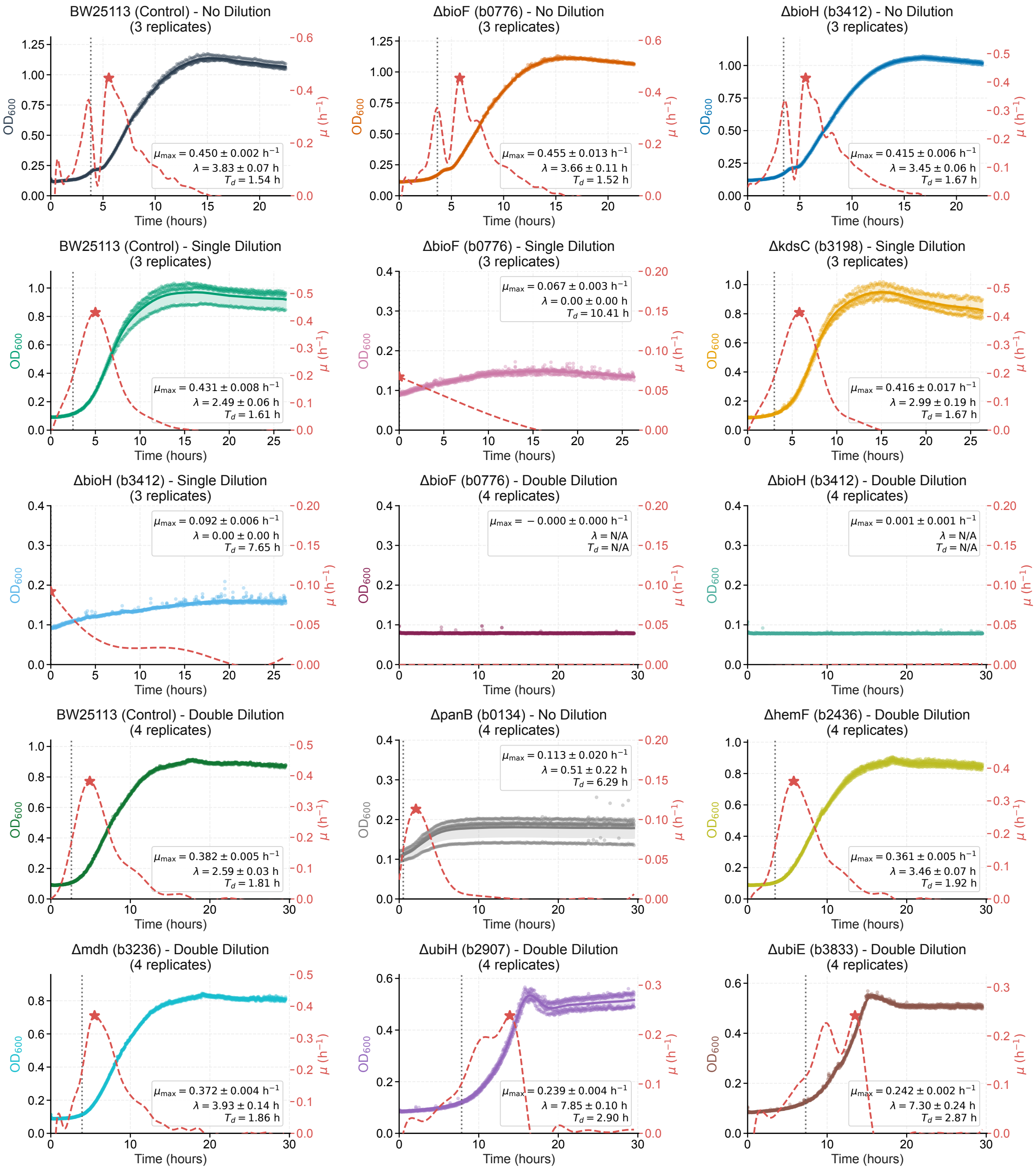


**Fig. B2**: **Kinetic growth profiling and `fitderiv` derivative analysis of *E. coli* deletion strains.** Cultivation profiles and growth rate derivative estimations across the indicated *E. coli* single-gene knockouts and parental BW25113 control strains under varying dilution regimes. Solid colored curves denote the Gaussian Process (GP) regression mean fit to empirical $\mathrm{OD}_{600}$ replicate measurements (individual points; shaded bands represent $\pm 1$ s.d.; $n = 3$–$4$ replicates per condition). Red dashed curves show the inferred instantaneous specific growth rate $\mu(t) = \frac{\mathrm{d}\ln(\mathrm{OD}_{600})}{\mathrm{d}t}$ ($\mathrm{h}^{-1}$) on the secondary (right) axis, with star markers indicating peak growth rates ($\mu_{\max}$). Inset boxes summarize the extracted kinetic parameters ($\mu_{\max}$, lag phase duration $\lambda$, and doubling time $T_d = \frac{\ln 2}{\mu_{\max}}$) reported as mean $\pm$ s.d.; conditions without detectable growth are denoted as N/A.

The attribution $\phi_i(x)$ for an individual feature $i$ represents its expected marginal contribution across all possible feature subsets (coalitions) $S \subseteq F \setminus \{i\}$:

$$\phi_i(x) = \sum_{S \subseteq F \setminus \{i\}} \frac{|S|!(|F| - |S| - 1)!}{|F|!} \Big[ f_x(S \cup \{i\}) - f_x(S) \Big], \tag{C4}$$

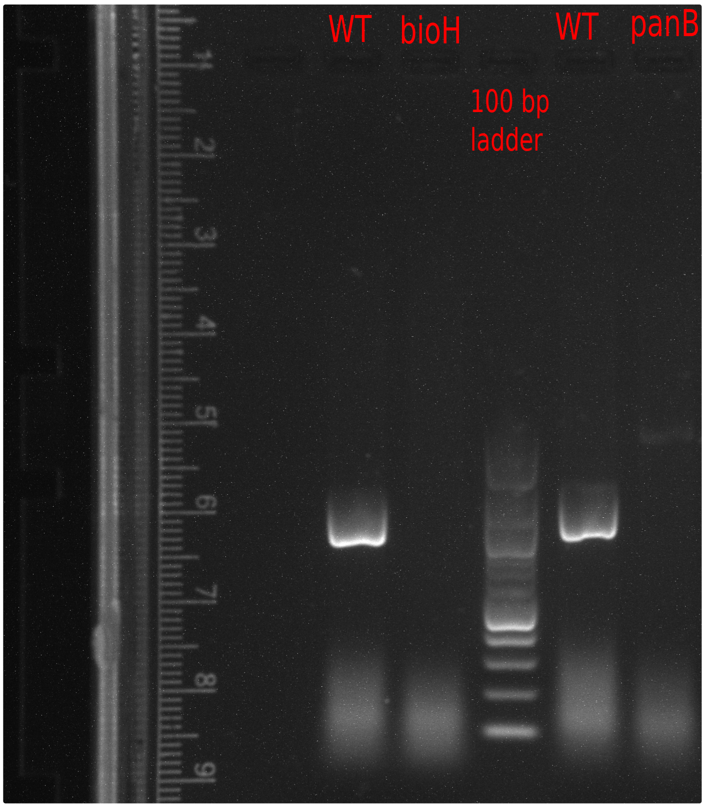


**Fig. B3**: **PCR validation of *bioH* and *panB* knockout strains.** Agarose gel electrophoresis of diagnostic PCR amplicons from parental wild-type *E. coli* BW25113 and target deletion mutants. Wild-type lanes yield expected bands at 973 bp (*bioH*) and 1020 bp (*panB*). The $\Delta$*panB* lane shows a size shift to a faint $\sim$1.3 kb band, corresponding to insertion of the kanamycin resistance cassette ($kan^R$), while no amplicon is detected for $\Delta$*bioH*. DNA ladder (100 bp) is shown for molecular size reference.

where $F$ denotes the complete set of features, $S$ is a subset excluding feature $i$, $f_x(S) = \mathbb{E}[f(X) \mid X_S = x_S]$ is the expected model prediction conditioned only on the feature subset $S$ (with unselected features marginalized out over a reference distribution), and the combinatorial factor $\frac{|S|!(|F|-|S|-1)!}{|F|!}$ represents the probability of coalition $S$ forming under a uniformly random feature permutation.

#### C.1 Kernel SHAP

To compute local feature attributions for our hypergraph model, we employ Kernel SHAP [5]. We formulate the explanation task by assigning each of the $M = |F| = 2712$ reactions a binary inclusion indicator: an active reaction (1) retains its original edge feature values, whereas an inactive or knocked-out reaction (0) has its corresponding edge features set to zero.

Because computing exact Shapley values across $2^{2712}$ possible reaction coalitions is computationally intractable, Kernel SHAP approximates these attributions by solving a weighted linear regression problem over a tractable subset of sampled coalitions. Specifically, for an input hypergraph characterized by its reaction feature matrix $\mathbf{X} \in \mathbb{R}^{2712\times 4}$ (denoted simply as $x$), Kernel SHAP executes the following steps:

1. **Coalition sampling:** Binary coalition vectors $z' \in \{0,1\}^M$ are sampled, where $z'_j = 1$ denotes that reaction $j$ is active and $z'_j = 0$ denotes that it is inactive.
2. **Mapping to input space ($h_x(z')$):** In standard tabular domains, omitted features are typically marginalized over background empirical distributions. In our hypergraph setting, an inactive reaction is explicitly represented by zero-masking its associated edge feature tensor:

$$[h_x(z')]_j = \begin{cases} x_j, & \text{if } z'_j = 1 \quad \text{(reaction active)}, \\ \mathbf{0}, & \text{if } z'_j = 0 \quad \text{(reaction inactive/knocked out)}, \end{cases} \tag{C5}$$

   where $x_j \in \mathbb{R}^4$ is the observed edge feature vector of reaction $j$ in the query hypergraph.

3. **Shapley kernel weighting:** Each binary coalition $z'$ is assigned a weight via the Shapley kernel $\pi_{x'}(z')$:

$$\pi_{x'}(z') = \frac{M-1}{\binom{M}{|z'|}|z'|(M-|z'|)}, \tag{C6}$$

where $|z'| = \sum_{j=1}^{M} z'_j$ is the number of active reactions in the coalition. This kernel assigns the highest weights to near-empty ($|z'| \approx 1$) and near-full ($|z'| \approx M-1$) coalitions, effectively isolating the marginal effects of individual reactions.

4. **Weighted least squares optimization:** A linear explanation surrogate $g(z') = \phi_0 + \sum_{j=1}^{M} \phi_j z'_j$ is fitted by minimizing the weighted squared loss:

$$\mathcal{L}(f, g, \pi_{x'}) = \sum_{z' \in \mathcal{Z}} \Big[ f\big(h_x(z')\big) - g(z') \Big]^2 \pi_{x'}(z'), \tag{C7}$$

where $\mathcal{Z}$ denotes the set of sampled coalitions, $\phi_0 = f(\mathbf{0})$ corresponds to the base prediction under a fully zero-masked hypergraph, and the resulting coefficients $\phi_j$ serve as the estimated SHAP values for each reaction $j$.

#### C.2 Gradient SHAP

Gradient SHAP (also known as Expected Gradients) [5, 6] extends the Integrated Gradients framework [7] to approximate Shapley values for differentiable neural architectures by integrating gradients along trajectories connecting the query input to a distribution of baseline reference points. By leveraging GPU-accelerated automatic differentiation, it calculates attributions across all input dimensions simultaneously in parallel backward passes. This circumvents the black-box formulation of Kernel SHAP, which requires evaluating thousands of discrete combinatorial feature coalitions, thereby avoiding the substantial computational overhead and prolonged execution times incurred by repeated forward model evaluations.

##### C.2.1 Baseline reference definition

The background baseline hypergraph $\mathbf{X}' \in \mathbb{R}^{2712 \times 4}$ was constructed by computing the feature-wise mean across all hypergraph reactions outside the test set. Each hyperedge (reaction) in $\mathbf{X}'$ was assigned this identical four-dimensional attribute vector: FBA $= 0.0799$, $\text{FVA}_{\text{max}} = 0.9796$, $\text{FVA}_{\text{min}} = -0.2782$, and pFBA $= 0.0800$. The query instance $\mathbf{X}$ corresponds to the intact sample hypergraph retaining its sample-specific hyperedge features.

##### C.2.2 Path-integral attribution

Gradient SHAP approximates the expected path gradient between the background baseline hypergraph $\mathbf{X}'$ and the query sample hypergraph $\mathbf{X}$ by sampling along straight-line interpolation trajectories. To mitigate high-frequency gradient noise and saturation artifacts arising from non-linear layer activations, a local smoothing perturbation $\boldsymbol{\epsilon} \sim \mathcal{N}(\mathbf{0}, \sigma^2\mathbf{I})$ with $\sigma = 0.1$ is injected at each step, yielding the perturbed interpolated hypergraph:

$$\widetilde{\mathbf{X}}_\alpha = \mathbf{X}' + \alpha(\mathbf{X} - \mathbf{X}') + \boldsymbol{\epsilon}, \quad \alpha \sim \mathcal{U}(0,1), \tag{C8}$$

where $\alpha$ denotes the uniform interpolation step parameter.

The attribution $\phi_{j,d}$ for the $d$-th feature dimension ($d \in \{1, \dots, 4\}$) of reaction hyperedge $j$ is then computed as:

$$\phi_{j,d}(\mathbf{X}) = (x_{j,d} - x'_{j,d}) \cdot \mathbb{E}_{\alpha \sim \mathcal{U}(0,1),\, \boldsymbol{\epsilon} \sim \mathcal{N}(\mathbf{0}, \sigma^2\mathbf{I})} \left[ \frac{\partial f(\widetilde{\mathbf{X}}_\alpha)}{\partial x_{j,d}} \right], \tag{C9}$$

where $x_{j,d}$ and $x'_{j,d}$ denote the observed and baseline feature values from $\mathbf{X}$ and $\mathbf{X}'$, respectively, and $\frac{\partial f(\widetilde{\mathbf{X}}_\alpha)}{\partial x_{j,d}}$ is the partial derivative of the HyperGenie model output $f$ evaluated at the perturbed state $\widetilde{\mathbf{X}}_\alpha$ via backpropagation.

##### C.2.3 Reaction-level importance aggregation

To quantify the overall importance of the $j$-th reaction hyperedge irrespective of attribution sign, we compute the total attribution magnitude by summing the absolute values of its four constituent feature attributions:

$$\Phi_j(\mathbf{X}) = \sum_{d=1}^{4} \left|\phi_{j,d}(\mathbf{X})\right|. \tag{C10}$$

The resulting non-negative scalar $\Phi_j(\mathbf{X}) \geq 0$ reflects the absolute contribution and structural sensitivity of reaction $j$ to the HyperGenie model prediction, preventing opposing positive and negative feature attributions from cancelling each other out.

### Appendix D Additional figures and tables

Figure D1 presents five precision-recall curves generated using HyperGenie. These curves correspond to one complete cycle of cross-validation, ensuring that each reaction appears in the test set exactly once. Figures D2A and D2B highlight two outlier runs with notably low PR-AUC values; these plots annotate five non-essential genes (according to the iML1515 experimental dataset) that were assigned high essentiality probabilities, thereby driving the drop in area under the curve.

Table D1 details the modified reaction bounds used to adapt the iML1515 metabolic model from the K-12 MG1655 strain to the BW25113 strain, for which experimental measurements are available. The table also specifies the reaction bounds (uptake rates) defining the nutrient conditions.

Table D2 shows the details of the hyperparameter grid search used to select the hyperparameters for the Hypergraph Neural Network. The bold values in the table indicate the chosen optimal hyperparameters, used to generate the results presented in this work.

Table D3 details the cross-condition predictive performance of HyperGenie when evaluating differentially essential genes, defined as genes essential in a target carbon source but dispensable in glucose. In this evaluation mode, the model was trained exclusively on essentiality data from glucose minimal medium (M9) and tasked with predicting condition-specific essentiality across eleven alternative nutrient conditions.

Finally, Figure D3 compares the PR-AUC performance of HyperGenie, HyperGenie with log-modulus normalized edge features (HyperGenie Norm), and MLPGenie across twelve carbon sources and three nutrient uptake rates (10, 20, and 40 mmol $\cdot$ gDW$^{-1}$ $\cdot$ h$^{-1}$). Performance notably improved for acetate, pyruvate, alanine, and D-lactate as the uptake rate increased from 10 to 20 and 40 mmol $\cdot$ gDW$^{-1}$ $\cdot$ h$^{-1}$, a trend that was especially pronounced for HyperGenie.

**Table D1**: **Reaction bounds for the *E. coli* iML1515 metabolic model.** The specific flux bounds required to adapt the standard K-12 MG1655-based iML1515 model to the BW25113 strain and simulate aerobic growth in glucose minimal media. Flux bounds are given in standard units of mmol gDW$^{-1}$ h$^{-1}$.

| **Reaction (BiGG ID)** | **Lower bound** | **Upper bound** |
|---|---|---|
| *Strain adaptation (K-12 MG1655 to BW25113)* | | |
| $\beta$-galactosidase (LACZ) | 0 | 0 |
| Rhamnulokinase (RMK) | 0 | 0 |
| L-rhamnose isomerase (RMI) | 0 | 0 |
| Lyxose isomerase (LYXI) | 0 | 0 |
| Rhamnulose-1-phosphate aldolase (RMPA) | 0 | 0 |
| L-ribulokinase (RBK_L1) | 0 | 0 |
| L-arabinose isomerase (ARAI) | 0 | 0 |
| *Aerobic growth conditions (minimal media)* | | |
| Oxygen exchange (EX_o2_e) | −20 | 1000 |
| Carbon source (e.g., D-glucose exchange, EX_glc__D_e) | −10 | 1000 |

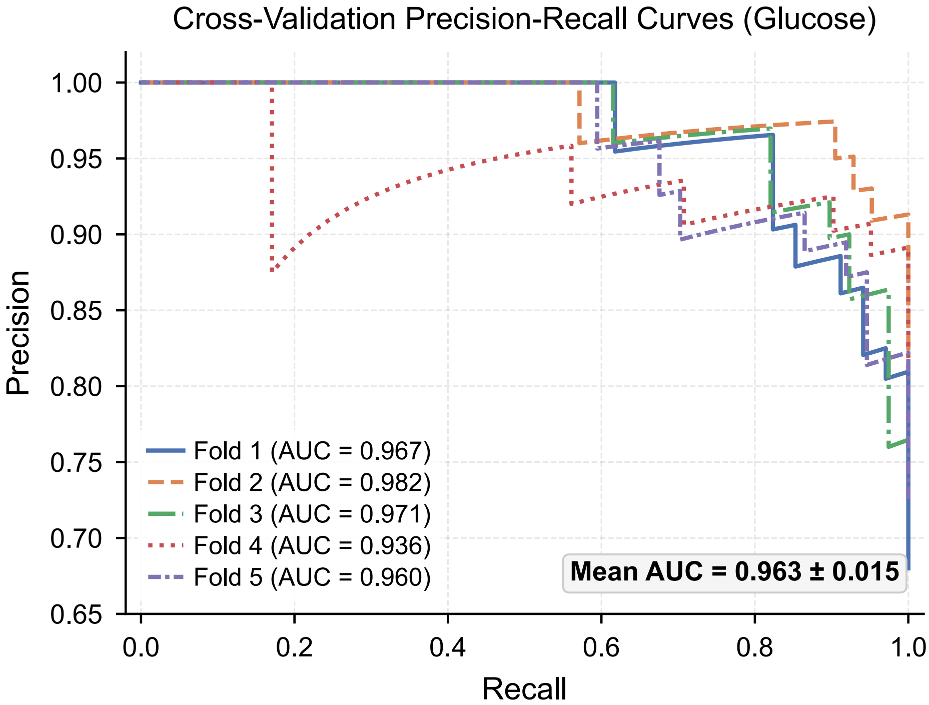


**Fig. D1**: **Precision-Recall curves are shown for five models**, each trained on a different 80% subset of the data, ensuring each data point is tested exactly once. This represents one cycle of a 5-fold cross-validation process for bacteria grown aerobically in glucose.

**Table D2**: **Hyperparameter optimization for HyperGenie and MLPGenie.** The evaluated search space and selected optimal values for the hypergraph neural network (HyperGenie) and the multilayer perceptron architecture (MLPGenie). The architecture and training parameters of MLPGenie structurally mirror the HyperGenie classification module, with the sole exception of the weight decay as explicitly noted. Parameters without a defined search space were held fixed during the evaluation.

| **Parameter** | **Search space** | **Optimal value** |
|---|---|---|
| *Architecture* | | |
| Hidden layer dimension ($MLP_{1,2,3,4}$) | {32, 64, 128, 256} | 64 |
| Classification layer dimension | {32, 64, 128, 256} | 128 |
| Number of HGNN layers | – | 2 |
| Number of fully connected layers | – | 2 |
| *Training* | | |
| Learning rate | $\{10^{-2}, 10^{-3}, 10^{-4}\}$ | $10^{-3}$ |
| Weight decay (HyperGenie) | {0.0, 0.01, 0.05} | 0.05 |
| Weight decay (MLPGenie) | {0.0, 0.01, 0.05} | 0.0 |
| Maximum epochs | – | 1600 |
| Dropout rate | – | 0.50 |

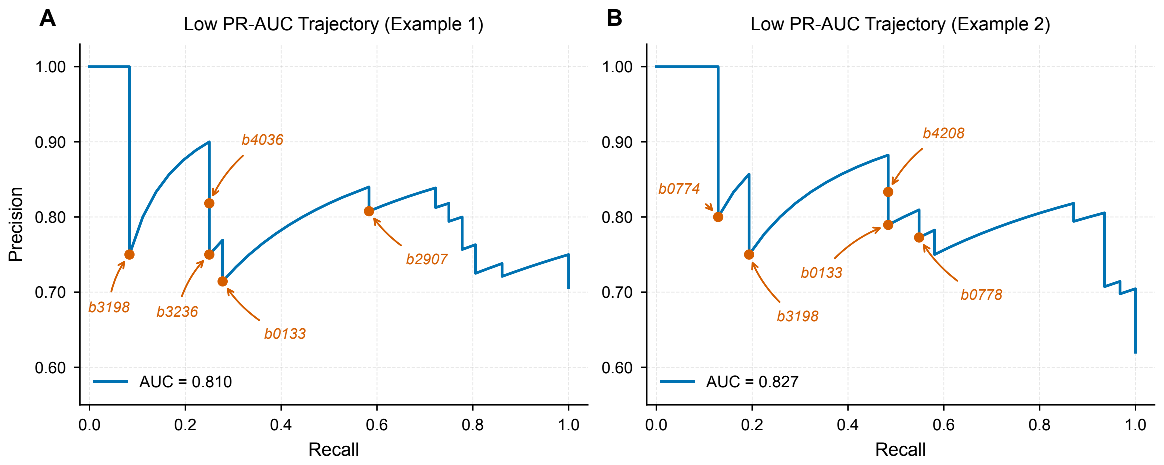


**Fig. D2**: **Error analysis of low-performing cross-validation folds.** (**A**, **B**) Precision–recall curves for two representative outlier folds exhibiting markedly low PR-AUC values. Labeled data points highlight non-essential genes according to the *i*ML1515 experimental dataset (*e.g.*, *b0133*, *b0774*, *b3198*, and *b3236*) that were assigned high essentiality probabilities by HyperGenie, representing prominent false positives.

**Table D3**: Cross-condition essentiality prediction. The model was trained on glucose essentiality data and evaluated on alternative nutrients. For each target nutrient, the table details the total number of expected differentially essential genes (essential in the target nutrient but dispensable in glucose), the specific genes correctly identified by the model, and the specific genes it failed to identify (false negatives).

| **Target Nutrient** | **Expected Genes** | **Correctly Predicted Genes** | **Missed Genes** |
|---|---|---|---|
| Acetate | 11 | *apaH*, *acnB*, *ubiF*, *thiD*, *ubiH*, *cyaA*, *ubiE* | *lipB*, *pta*, *mdh*, *aceA* |
| Pyruvate | 11 | *apaH*, *ubiF*, *aspC*, *thiD*, *fruK*, *ubiH*, *rpe*, *ubiE* | *lipB*, *ppsA*, *avtA* |
| Succinate | 7 | *ubiF*, *thiD*, *ubiH*, *cyaA*, *ubiE* | *lipB*, *dctA* |
| Alanine | 8 | *lipB*, *ubiF*, *thiD*, *ubiH*, *rpe*, *cyaA*, *ubiE* | *acpH* |
| Oxoglutarate | 8 | *ubiF*, *thiD*, *kgtP*, *ubiH*, *hldE*, *cyaA*, *ubiE* | *lipB* |
| Xylose | 13 | *panC*, *panB*, *ubiF*, *gnd*, *thiD*, *ubiH*, *rpe*, *xylB*, *cyaA*, *ubiE* | *ldcA*, *pgpB*, *xylA* |
| Sorbitol | 9 | *panC*, *panB*, *thiD*, *ubiH*, *mtlD*, *cyaA*, *ubiE* | *purU*, *rpe* |
| Ribose | 7 | *thiD*, *ubiH*, *rpe*, *rbsK*, *cyaA*, *ubiE* | *lipB* |
| Gluconate | 7 | *speE*, *thiD*, *ubiH*, *rpe*, *cyaA*, *ubiE* | *eda* |
| Galactose | 7 | *galK*, *galT*, *galE*, *thiD*, *ubiH*, *cyaA*, *ubiE* | – |
| D-Lactate | 8 | *ubiF*, *thiD*, *fruK*, *ubiH*, *cyaA*, *ubiE* | *lipB*, *ppsA* |

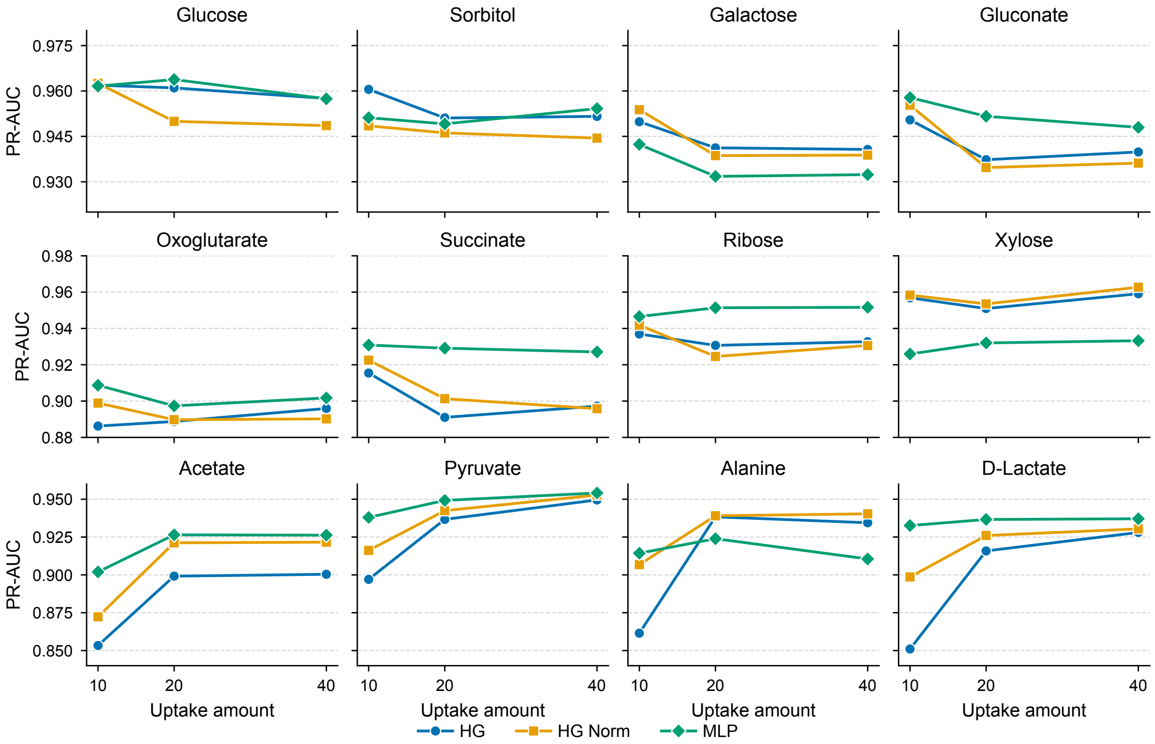


**Fig. D3**: **Impact of nutrient uptake rates on model predictive performance across carbon sources.** PR-AUC scores for HyperGenie, HyperGenie Norm, and MLPGenie evaluated across twelve carbon sources in minimal media at three uptake rates: 10, 20, and 40 $\text{mmol} \cdot \text{gDW}^{-1} \cdot \text{h}^{-1}$. Substantial performance gains are observed for acetate, pyruvate, alanine, and D-lactate when increasing uptake rates from 10 to 20 and 40 $\text{mmol} \cdot \text{gDW}^{-1} \cdot \text{h}^{-1}$, with HyperGenie demonstrating the most substantial improvements.

#### References


[1] Monk, J.M., Lloyd, C.J., Brunk, E., Mih, N., Sastry, A., King, Z., Takeuchi, R., Nomura, W., Zhang, Z., Mori, H., *et al.*: iML1515, a knowledgebase that computes Escherichia coli traits. Nature biotechnology **35**(10), 904–908 (2017)

[2] Orth, J.D., Thiele, I., Palsson, B.Ø.: What is flux balance analysis? Nature biotechnology **28**(3), 245–248 (2010)

[3] Swain, P.S., Stevenson, K., Leary, A., Montano-Gutierrez, L.F., Clark, I.B., Vogel, J., Pilizota, T.: Inferring time derivatives including cell growth rates using Gaussian processes. Nature communications **7**(1), 13766 (2016)

[4] Shapley, L.: A Value for $n$-Person Games. In: Kuhn, H., Tucker, A. (eds.) Contributions to the Theory of Games, Volume II, pp. 307–317. Princeton University Press, Princeton (1953)

[5] Lundberg, S.M., Lee, S.: A unified approach to interpreting model predictions. Advances in neural information processing systems **30** (2017)

[6] Erion, G., Janizek, J.D., Sturmfels, P., Lundberg, S.M., Lee, S.: Improving performance of deep learning models with axiomatic attribution priors and expected gradients. Nature machine intelligence **3**(7), 620–631 (2021)

[7] Sundararajan, M., Taly, A., Yan, Q.: Axiomatic attribution for deep networks. In: International Conference on Machine Learning, pp. 3319–3328 (2017). PMLR
